# KLF5 controls subtype-independent highly interactive enhancers in pancreatic cancer to regulate cell survival

**DOI:** 10.1101/2025.06.30.662301

**Authors:** Thomas L Ekstrom, Zhangshuai Dai, Julia Thiel, Meghana Manjunath, Nadine Schacherer, Bishakha Joyeeta Saha, Yara Souto, Amro M Abdelrahman, Frank Essmann, Sven Beyes, Mark J Truty, Meng Dong, Steven A Johnsen

## Abstract

Pancreatic ductal adenocarcinoma (PDAC) remains a highly lethal cancer with a 5-year survival rate of 13%. Despite recent molecular stratification of tumors into distinct classical and basal-like cell states, most tumors are heterogeneous contain of both subtypes. Therefore, therapeutic approaches targeting one subtype may not be suitable for PDAC therapy. Here, we integrated chromatin accessibility (ATAC-seq), genome-wide occupancy (ChIP-seq) for epigenetic status (H3K27ac) and H3K4me3-anchored chromatin topology (HiChIP) to uncover subtype-independent highly interactive enhancers that interact with essential genes in PDAC. Motif analysis revealed these common enhancers were bound by KLF5 with subsequent depletion leading to decreased cell viability via induction of apoptosis. To elucidate the transcriptional and epigenetic mechanisms by which KLF5 functions in PDAC, we employed rapid depletion of KLF5 with dTAG technology and profiled the effects on the open and active chromatin landscape and transcription with nascent RNA and mRNA-seq over time. Enhancer inactivation via KRAB domain Zim3-dCas9 fusion protein confirmed KLF5-bound enhancers regulate target genes, including the anti-apoptotic gene *BCL2L1.* Multiplex immunofluorescence confirmed co-staining of KLF5 and Bcl-xL in patient samples and overexpression of Bcl-xL rescued the induction of apoptosis after KLF5 depletion. Taken together, this study provides new insights into common mechanisms to target highly heterogeneous PDAC tumors.

**Teaser:** KLF5 controls subtype-independent highly interactive enhancers to regulate cell viability in pancreatic cancer.

## Introduction

Pancreatic ductal adenocarcinoma (PDAC) remains a lethal cancer with a 5-year survival rate of 13% (*1*). Recent molecular stratification of patient tumors into distinct classical and basal-like cell states have revealed subtype-specific responses to therapy (*2*). However, PDAC tumors are highly heterogeneous usually containing cells of each subtype as well as cells in intermediate co-expressing states (*3*). Thus, therapeutic approaches targeting one subtype may not be suitable for many patients. In this study, we sought to identify subtype-independent transcription factor-mediated epigenetic and transcriptional regulatory mechanisms that govern cell survival in pancreatic cancer.

The Krüppel-like factor (KLF) family of zinc-finger transcription factors consisting of 17 members. Under normal conditions, KLF5 regulates differentiation and development and its deletion is embryonic lethal in mice (*4*). In pancreatic cancer, KLF5 is overexpressed and its conditional deletion reduces acinar to ductal metaplasia (ADM) and pancreatic intraepithelial neoplasia (PanIN) lesion formation in the Kras^LSLG12D/+;^ Ptf1a-Cre^ERTM^ PDAC model, suggesting KLF5 plays a role in PDAC pathobiology (*5*). Additional studies have interrogated CRISPR-dependency data and found gastrointestinal, squamous, ovarian and pancreatic cancers are among the most dependent on KLF5 (*6*). Mechanistically, KLF5 binds enhancers in promoter hubs (many enhancers per promoter (*7*)) and interacts with p63 and CBP in squamous carcinomas, ultimately increasing histone acetylation at enhancers to promote target gene expression via recruitment of BRD4 leading to Pol II pause release (*6*). Consistently, KLF5 was shown to maintain the active epigenetic landscape in PDAC by binding enhancer regions (*8*). KLF5 has also been shown to be overexpressed in cancers through amplification of its super enhancer, increased KLF5 protein stability through missense mutations that disrupt KLF5-FBXW7 interactions and mutations in the zinc-finger domain which alter its DNA-binding specificity (*9*) (*10*). Taken together, these data point to KLF5 as a putative target in pancreatic cancer. To this end, a small molecule inhibitor, ML264, has been shown to selectively decrease KLF5 expression and restore sensitivity in colorectal cancer patient-derived organoids (*11*). However, due to the intrinsic disordered regions (IDR) of KLF5, targeting the activity of KLF5 remains challenging. Thus, finding the molecular pathways and downstream effectors that KLF5 acts upon will provide a basis for identifying new therapeutic targets for pancreatic cancer.

In this study, we probed genome-wide CRISPR-Cas9 knockout screening data to identify subtype-independent mechanisms of cell survival in pancreatic cancer. We filtered top hits for transcription factors that bind subtype-independent highly interactive enhancers. To this end, we identified KLF5 to be enriched at these loci and depletion of KLF5 decreased cellular viability via induction of apoptosis. We further investigated the mechanistic underpinnings of these effects using an endogenous knock-in cell line of GFP-FKBP12^F36V^ into the N-terminus of *KLF5* to induce rapid degradation, thereby enabling us to monitor the primary transcriptional targets and regulatory mechanisms. We found degradation of KLF5 at early time points preferentially leads to decreased expression target genes connected with multiple KLF5-bound loci. Consistently, siRNA-mediated depletion of KLF5 across multiple cell lines showed KLF5-dependent genes interact with multiple KLF5-bound enhancers compared to KLF5-independent genes. This suggests KLF5 functions through hubs to regulate its target genes. A prominent example is *BCL2L1*, which encodes the anti-apoptotic protein Bcl-xL. Multispectral staining of patient tumors showed co-staining and enhancer inactivation via KRAB domain Zim3-dCas9 fusion protein confirmed KLF5-bound enhancers promote *BCL2L1* expression, thereby limiting apoptosis induction. Taken together, this study uncovers new insights into the molecular underpinnings of subtype-independent survival mechanisms in pancreatic cancer.

## Results

### Identification of subtype-independent TF-mediated highly interactive enhancers

To discern subtype-independent mechanisms of cell survival in pancreatic cancer, we probed publicly available genome-wide CRISPR knockout screen data from the Broad Institute DepMap project. Recently, specific transcription factors have been shown to regulate enhancer hubs to promote oncogenic pathways in cancer (*12–15*). Thus, we hypothesized that subtype-independently expressed transcription factors may regulate highly interactive enhancers in pancreatic cancer irrespective of molecular identity. To this end, we utilized our previously published assay for transposase-accessible chromatin with sequencing (ATAC-seq) to identify accessible chromatin regions in both classical (AsPC1) and basal-like (L3.6pl) PDAC cells (*16*). We subtracted the transcriptional start site (TSS) to gain distal open regions in each cell line and intersected these regions with H3K27ac peaks for distal open active (DOA) regions. We further subtracted CTCF peaks in each respective cell line to garner transcription factor-mediated DOA regions that were not simply structural interactions (i.e., TAD-boundaries). We then intersected these regions in AsPC1 with L3.6pl and identified 3993 subtype-independent TF-mediated DOA regions. Utilizing our H3K4me3-anchored HiChIP data in each cell line, we filtered regions that interact with our subtype-independent TF-mediated DOA regions to an active TSS marked with H3K4me3 and further filtered for enhancers with greater than or equal to 5 interactions to a specific gene. This number was chosen based as the inflection point of interaction distribution in each cell line (fig. S1A). This workflow resulted in 1310 regions that are marked by H3K27ac in multiple classical (AsPC1, HPAFII, TCCPAN2) and basal-like (BxPC3, L3.6pl, T3M4) cell lines, further confirming their subtype independence (fig. S1B).

To assess which TFs may play a role in regulating these enhancer-promoter integrations (EPI), we further interrogated these 1310 regions. Briefly, we performed motif analysis and filtered the CRISPR-Cas9 screen for dependent genes and found the KLF5 motif to be enriched at these loci (Fig. 1 A and B and fig. S1C). Interestingly, nine out of ten top motifs belonged to the activator-protein 1 (AP1) family of transcription factors with only JUNB found to influence viability in both subtypes (average gene effect of −0.51 and −0.49 in classical and basal-like cells, respectively) (fig. S1C). To validate these *in* silico findings, we performed KLF5 ChIP-seq in four different PDAC cell lines that were shown to be dependent on KLF5 in DepMap data (Fig. 1C and fig. S1D), AsPC1, HPAFII, BxPC3 and L3.6pl, and show KLF5 binding at these enhancers in all cell lines, confirming our bioinformatic analyses (Fig. 1C). Consistent with the reported role of KLF5 at enhancers, genomic annotation analyses showed KLF5 primarily binds distal regions (fig. S2A). We observed the strongest binding and motif enrichment of KLF5 at common or mixed (present in two or more but not all cell lines) regions (fig. S2 B and C). Lastly, aggregate peak analysis (APA) validated the strength of interactions that originate from these subtype-independent TF-mediated DOA regions (Fig. 1D).

**Fig. 1.**
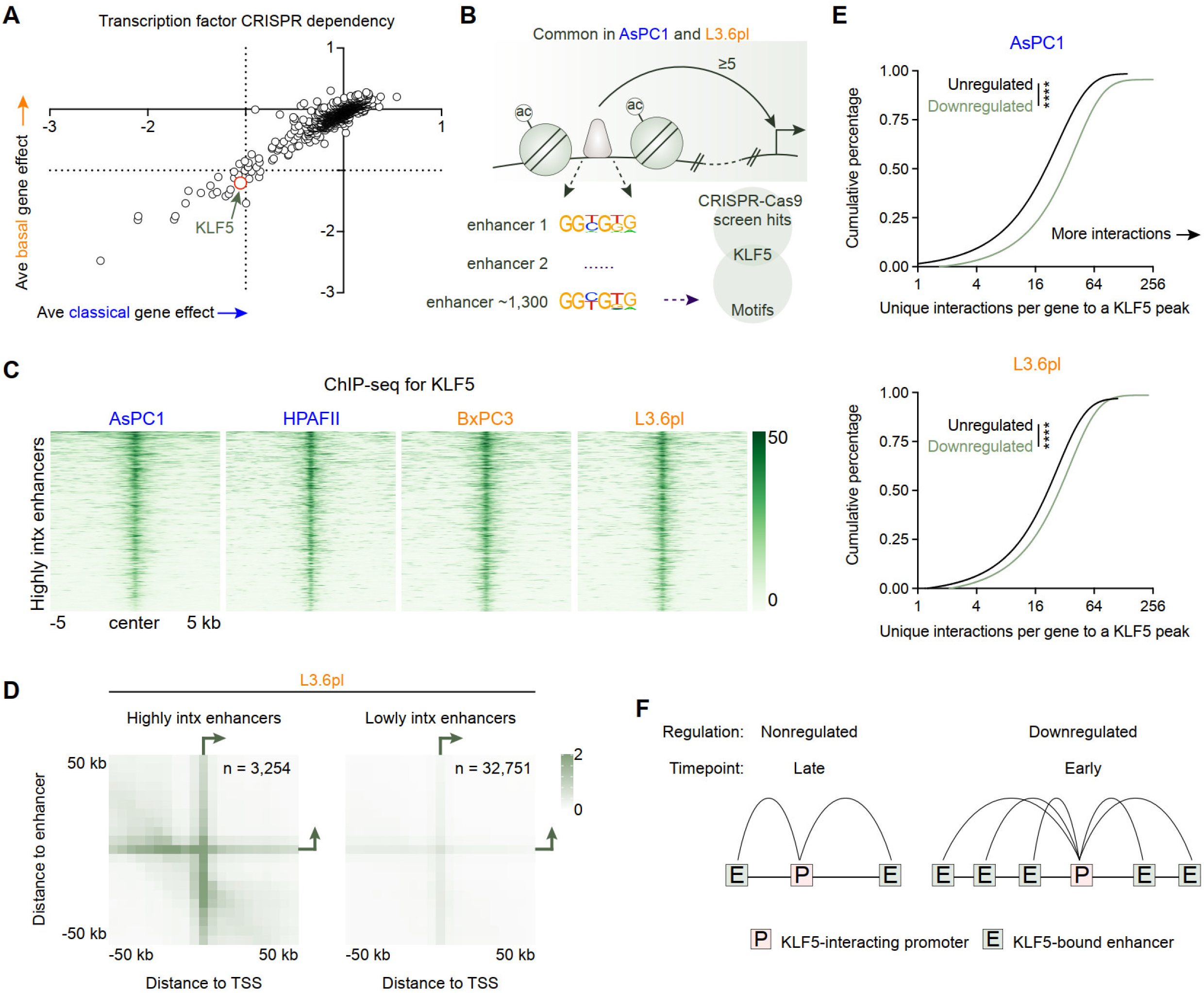
KLF5 regulates pancreatic cancer cell viability in a subtype-independent manner. (**A**) CRISPR-dependency data from DepMap for transcription factors averaged over classical (AsPC1, HPAFII, PaTu8988s, SUIT2, TCCPAN2) and basal-like A (BxPC3, DanG, Hs766t, L3.3, PK1, T3M4) cell lines. (**B**) Schematic depicting workflow of identifying subtype-independent highly interactive enhancers. The motifs from these regions were intersected with the dependent genes from (A). (**C**) Heatmap of KLF5 read density in AsPC1, HPAFII, BxPC3 and L3.6pl cells at subtype-independent highly interactive enhancers. Reads normalized to reads per genome coverage (RPGC), n = 2 biological replicates. (**D**) Aggregate peak analysis of highly interactive (contact counts ≥ 5) and lowly interactive (contact counts < 5) enhancers in L3.6pl cells. Normalized to the number of loops (number of rows present in the bedpe file). (**E**) Cumulative percentage plots of the number of KLF5 peaks that interact with H3K4me3 marked KLF5-dependent genes (Log2FC < −0.25, FDR < 0.05) or KLF5-nonregulated (−0.15 < Log2FC < 0.15, FDR > 0.25) genes across all four (AsPC1, HPAFII, BxPC3, L3.6pl) cell lines in AsPC1 and L3.6pl cells. 355 down genes and 396 nonregulated were included in the analysis. Mann-Whitney test (Wilcoxon rank-sum test), **** p < 0.0001. (**F**) Schematic depicting the number of KLF5 peaks that connect to 1) KLF5-nonregulated and KLF5-dependent or 2) early and late KLF5-depdnent genes.

To discern the transcriptional effects elicited by KLF5 depletion, we performed mRNA-seq upon KLF5 knockdown (KD) in all four cell lines and found KLF5 primarily regulates gene sets associated with differentiation, developmental and oncogenic pathways (fig. S3 A and B). Interestingly, common KLF5-dependent genes across all four cell lines are more connected to multiple distal KLF5 binding sites (in AsPC1 and L3.6pl cells) compared to nonregulated genes (Fig. 1 E and F) as assessed by mRNA-seq (fig. S3C). Lastly, we observed moderate and highly mixed effects on subtype identity, with limited overlap between KLF5 occupancy and that of GATA6 or ΔNp63 in AsPC1 and L3.6pl, respectively (fig. S4 A and B). To confirm the dependency of PDAC on KLF5, we performed crystal violet staining upon KLF5 depletion in AsPC1, HPAFII, BxPC3 and L3.6pl cells (fig. S5 A to C). HPAFII, BxPC3 and L3.6pl cells showed a robust decrease in crystal violet staining, whereas AsPC1 cells showed a more moderate effect of KLF5 depletion on cell viability. Lastly, we show that KLF5 knockdown has no detectable effects on cell viability in the immortalized normal human pancreatic ductal epithelial cell line (HPDEC), supporting that KLF5 represents a cancer-specific dependency (fig. S5 A to C).

### Rapid degradation of KLF5 primarily affects distal enhancers

Because loss of KLF5 leads to decreased cell viability, we employed dTAG technology to rapidly degrade KLF5 and perform kinetic analyses following its perturbation. Briefly, we utilized CRISPR-Cas9 to knockin GFP-FKBP12^F26V^ into the N-terminus of the endogenous *KLF5* gene (Fig. 2A, referred to as knockin cells). We chose the N-terminus of *KLF5* as the C-terminus contains the zinc finger binding domains, whereas the N-terminus contains the intrinsic disordered regions (IDR). Western blot analysis shows near complete depletion of KLF5 protein levels after 1 h treatment of 250 nM VHL-based degrader dTAGV-1 TFA (referred to as dTAG) in a clonal homozygous L3.6pl cell line (Fig. 2B and fig. S6 A to C). We were able to generate homozygous knockin clones in AsPC1, BxPC3 and L3.6pl cells, but were unable to generate HPAFII knockin cells (fig. S6 A to C). We performed ChIP-seq for KLF5, H3K27ac and H3K4me3 as well as ATAC-seq upon early (1 h), intermediate (4 h) and late (24 h) time points after treatment with dTAG to assess the early and late KLF5-dependent epigenetic changes. ChIP-seq of KLF5 tagged with GFP-FKBP12^F26V^ in knockin L3.6pl cells showed similar genomic occupancy compared to wild-type cells (Fig. 2C) and dTAG treatment resulted in loss of KLF5 on chromatin (Fig. 2D) with more highly occupied KLF5 regions more sensitive to degradation (top of heatmap). We were able to detect differential levels of H3K27ac at KLF5 binding sites at early time points, but could identify regions with decreased H3K27ac at the intermediate and late time points (Fig. 2E). Interestingly, decreased H3K27ac occupancy at early and intermediate time points occurred primarily at distal regions, whereas the late time point showed changes primarily at promoters (Fig. 2F) and did not alter H3K4me3 levels (fig. S7A). In order to determine if these changes were caused by loss of chromatin accessibility, we performed ATAC-seq upon KLF5 degradation. Notably, we found that chromatin remained open at these enhancers following KLF5 depletion (Fig. 2G) with minimal overlap of differentially bound H3K27ac and ATAC levels (fig. S7 B and C). These results are consistent with prior reports of KLF5 interacting with CBP (*6*).

**Fig. 2.**
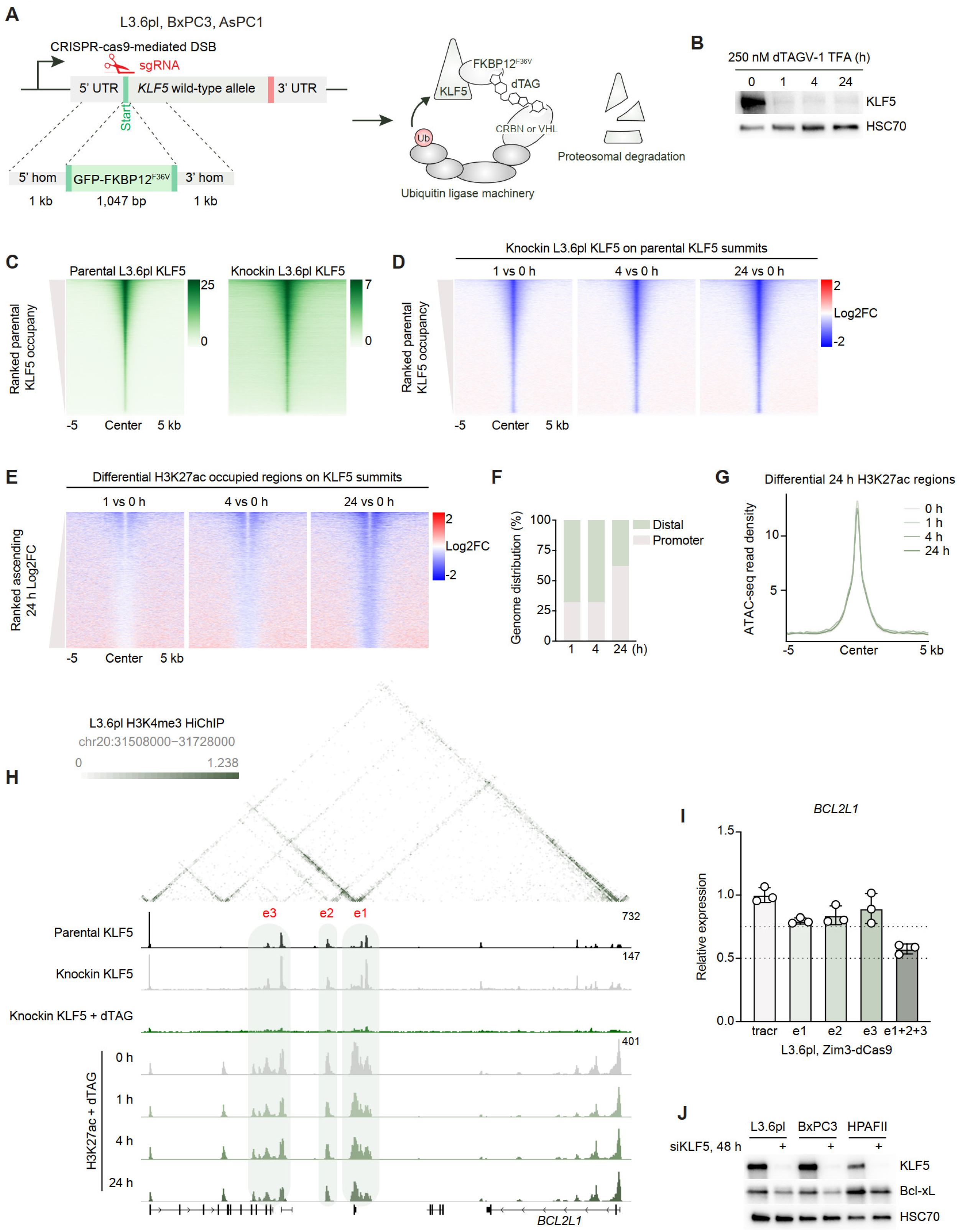
Rapid degradation of KLF5 preferentially targets distal regulatory elements. (**A**) Schematic depicting endogenous knock-in approach of GFP-FKBP12^F36V^ into the N-terminus of the *KLF5* locus with subsequent degradation. (**B**) Representative Western blot analysis of KLF5 after 250 nM dTAG treatment for 0, 1, 4 and 24 h. HSC70 was used as a loading control. (**C**) Heatmap of KLF5 read density of parental L3.6pl cells (left) and knockin L3.6pl cells (right) ranked on parental L3.6pl cells. Reads normalized to reads per genome coverage (RPGC), 85,262 regions were plotted, n = 2 biological replicates. (**D**) Log2FC heatmap of global KLF5 read density in knockin L3.6pl cells at 1, 4 and 24 h dTAG treatment ranked from regions in (C). Reads normalized to RPGC, n = 2 biological replicates. (**E**) Log2FC heatmap of differentially bound H3K27ac read density at KLF5 loci in knockin L3.6pl cells at 1, 4 and 24 h dTAG treatment ranked ascending. Reads normalized to RPGC, 6,719 KLF5 summits at 4,226 differentially bound H3K27ac regions were plotted, n = 3 biological replicates. (**F**) Genomic annotation of sites described in (E). (**G**) Aggregate plot of chromatin accessibility in knockin L3.6pl cells at 24 h dTAG treatment on regions in (E). (**H**) Contact matrix showing counts per million normalized contacts at the *BCL2L1* locus interacting with KLF5 binding sites that lose downstream distal H3K27ac sites at early timepoints. 2.5 kb bin size. Green shading highlights enhancers targeted for Zim3-dCas9-mediated repression. (**I**) qPCR of *BCL2L1* upon 48 h Zim3-dCas9 simultaneously targeting all three marked enhancers in L3.6pl cells. *ACTB* was used to normalize gene expression. n = 3 biological replicates. (**J**) Representative Western blot analysis of KLF5 and Bcl-xL in L3.6pl, BxPC3 and HPAFII cells after 48 h KLF5 knockdown. HSC70 was used as a loading control.

To explore the effects of KLF5 degradation on gene expression, we performed mRNA-seq 4, 24 and 48 h after dTAG treatment in knockin L3.6pl cells. Consistent with the maintenance of KLF5 binding observed in parental and knockin L3.6pl cells, siRNA- and dTAG-mediated depletion of KLF5 produced similar gene expression changes (fig. S8 A to C) and pathway regulation (fig. S8D). Taken together with the ChIP-seq, this suggests the addition of GFP-FKBP12^F26V^ does not alter the endogenous function of KLF5. Because we observed KLF5-dependent genes are connected to multiple distal KLF5 binding sites compared to nonregulated genes (Fig. 1D) and early enhancers were primarily distal (Fig. 2 E and F), we asked if early KLF5-dependent genes interacted with multiple distal KLF5 binding sites compared to late KLF5-dependent genes. In line with this hypothesis, we found genes downregulated after 4 h of KLF5 degradation connected with more KLF5 binding sites compared to later time points (fig. S8E).

### Enhancer inactivation of KLF5-bound enhancers regulates target gene expression

One example of an early KLF5-dependent gene that interacts with multiple distal KLF5 binding sites with decreased H3K27ac occupancy upon dTAG treatment and shown to be essential from the DepMap database is *BCL2L1*, which encodes the anti-apoptotic protein Bcl-xL. To discern the contribution of KLF5-bound enhancers to *BCL2L1* expression, we generated lentiviral Zim3-dCas9 L3.6pl and BxPC3 cell lines. We designed single guide RNAs (sgRNA) targeting enhancers that contain multiple KLF5 binding sites and display H3K27ac occupancy in all four cell lines as well as interacts with the TSS of *BCL2L1* (Fig. 2H and fig. S9A). Targeting these enhancers led to a reduced *BCL2L1* gene expression (Fig 2I and fig. S9B). We further confirmed that KLF5 depletion resulted in decreased Bcl-xL protein levels in L3.6pl, BxPC3 and HPAFII cell lines after siRNA-mediated (Fig. 2J). Repression of individual enhancer elements led to a weaker repression of *BCL2L1* gene expression compared to concomitantly targeting all elements (Fig 2I and fig. S9B). We suggest early KLF5-regulated genes such as *BCL2L1* are dependent on interconnected enhancer hubs that possess a multiple KLF5-bound regions required for gene activation and pancreatic cancer cell survival.

We next sought to identify if the enhancer of *BCL2L1* is transcribed in classical and basal-like cell lines and patient-derived xenografts (PDX). To this end, we analyzed our previously published precision run-on (PRO-seq) and length-extension chromatin run-on (leChRO-seq) data and found these loci are transcribed *in vitro* and *in vivo* in both PDAC subtypes (Fig. 3A) (*16*). Degradation of KLF5 in knockin L3.6pl, BxPC3 and AsPC1 cells showed downregulation of both *BCL2L1* enhancer (Fig. 3C) and pre-mRNA after 1 h dTAG treatment (Fig. 3B). In line with a downregulation of *BCL2L1* enhancer and pre-mRNA after KLF5 degradation, dTAG treatment of all three clonal knockin cell lines led to a decreased cell viability (fig. S9 C to E). Because we found KLF5 degradation influences *BCL2L1* eRNA transcription, we asked if KLF5 degradation influences global enhancer RNA transcription at KLF5-dependent loci. Thus, we performed 5-ethynyluridine (EU)-based nascent RNA profiling upon KLF5 degradation in knockin L3.6pl cells to identify rapid changes in RNA synthesis in proliferating cells. We focused on distal regions differentially occupied by H3K27ac upon dTAG treatment (Fig. 2E) and found that regions displaying rapid effects of KLF5 degradation on H3K27ac also displayed substantially higher levels of transcription, which decreased after KLF5 depletion, while enhancers affected at 4 24 h after dTAG treatment displayed decreased eRNA transcription and little (4 h) or no (24 h) effect of KLF5 degradation (Fig. 3D). To identify comparable control enhancer regions of interest, we identified distal accessible chromatin marked with H3K27ac and actively transcribed via detection of regulatory element (dREG) but devoid of KLF5. KLF5 degradation did not affect nascent RNA production at these control regions (Fig. 3D). We propose enhancer dependency on KLF5 is associated with increased eRNA transcription.

**Fig. 3.**
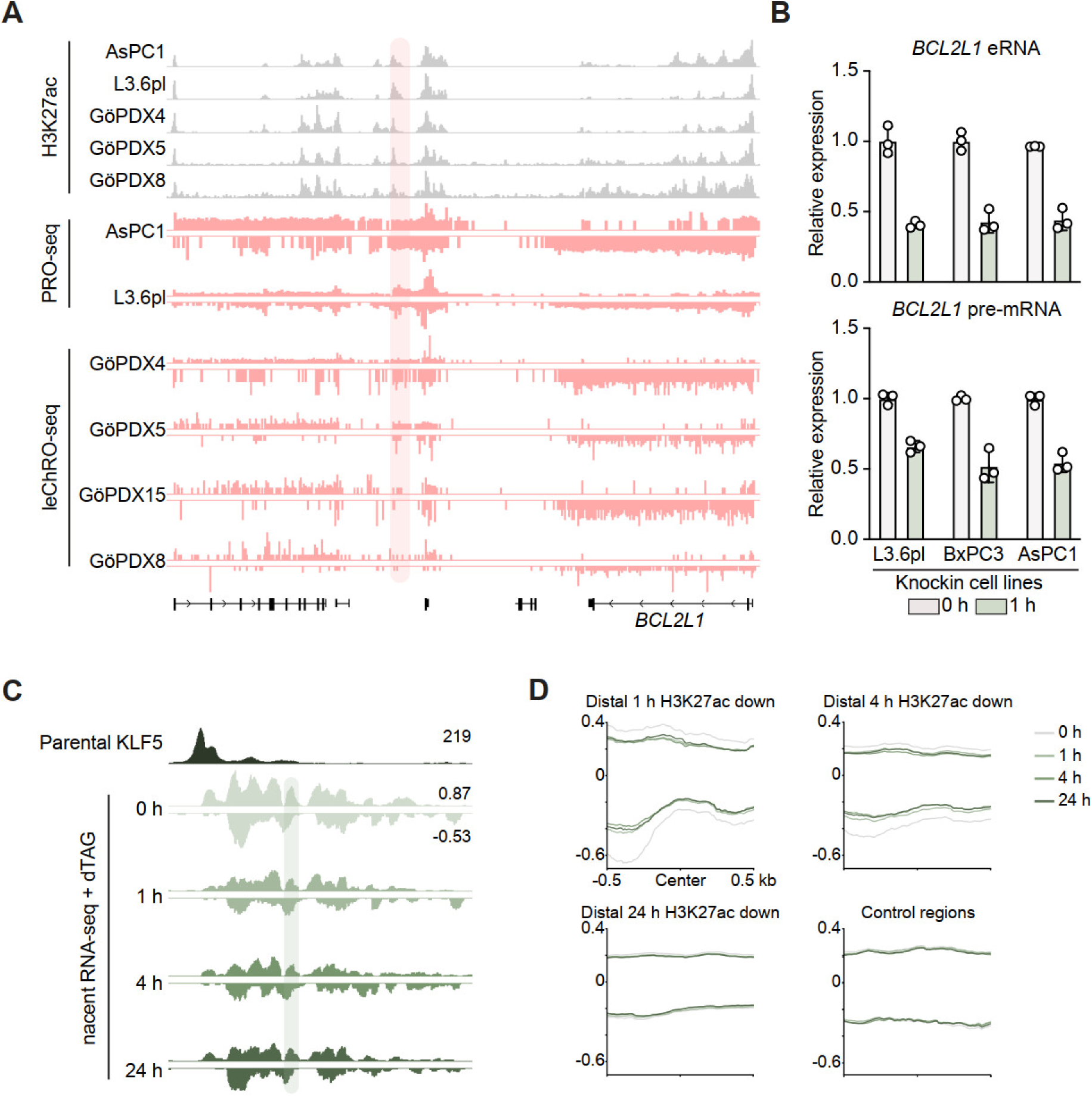
Enhancer RNA transcription is sensitive to early KLF5 degradation. (**A**) Integrated genome viewer (IGV) of H3K27ac ChIP-seq and nascent RNA profiling in cell lines (AsPC1, L3.6pl) and patient-derived xenografts (GöPDX4, GöPDX15, GöPDX5, GöPDX8) at the *BCL2L1* locus. Red shading highlights the site represented in (C). ChIP-seq reads normalized to reads per genome coverage (RPGC) and nascent RNA normalized to counts per million (cpm). (**B**) qPCR of *BCL2L1* eRNA (top) and premRNA (bottom) in knockin L3.6pl, BxPC3 and AsPC1 cells after 1 and 4 h dTAG treatment. *ACTB* was used to normalize gene expression. n = 3 biological replicates. (**C**) IGV track of parental L3.6pl cell KLF5 binding and EU-based nascent RNA-seq in knockin L3.6pl cells upon 0, 1, 4 and 24 h dTAG treatment downstream of the *BCL2L1* locus. Green shading marks the eRNA primer sites in (B). Reads normalized to cpm. (**D**) Aggregate plot of nascent RNA-seq in knockin L3.6pl cells after 0, 1, 4 and 24 h dTAG treatment on distal differential bound H3K27ac regions identified in (Fig. 2E). cpm normalized, 158, 406, 1,572, 516 regions were plotted for 1, 4, 24 h and control conditions, respectively, n = 3 biological replicates.

### KLF5 and Bcl-xL are co-expressed in PDAC patient tumors

To understand if the expression KLF5-dependent genes is, in fact, correlated with *KLF5* gene expression in patients we performed Pearson correlation on the cancer genome atlas (TCGA) pancreatic cancer dataset and found ∼60% of target genes, including *BCL2L1*, significantly and positively correlated with *KLF5* gene expression (Fig. 4A). Consistent with pancreatic cancer cell line dependency on KLF5, *KLF5*-hi/*BCL2L1*-hi patients showed a worse overall survival compared to *KLF5*-lo/*BCL2L1*-lo patients (Fig. 4A). We next sought to understand if KLF5-dependent genes such as *BCL2L1* are correlated with their dependency in cell lines. We conducted Pearson correlation on the publicly available genome-wide CRISPR-mediated gene knockout screen data and found *BCL2L1* to be among the top genes with correlated dependency with KLF5 (Fig. 4B). Because TCGA datasets are bulk-derived RNA-seq data, we asked if *KLF5* is correlated with dependent genes at single-cell resolution in patients. To this end, we merged single-cell RNA-seq (scRNA-seq) from six different publicly available datasets (*17–22*). We found *BCL2L1* and *KLF5* gene expression to be significantly correlated and expressed independent of the cell state (*3*) present in patients (Fig. 4 C and D and fig. S10A). To further confirm the co-expression of KLF5 and Bcl-xL in patients, we performed multiplex immunofluorescence (mIF) staining of four primary patient samples. Independent of subtype, we observed co-staining of KLF5 and Bcl-xL in both HNF4A-pos/KRT5/6-neg (classical) and HNF4A-neg/KRT5/6-pos (basal) cells within the same tumor (Fig. 4E and fig. S10B) and across all four patients (Fig. 4F), suggesting KLF5 acts in a subtype-independent manner to regulate Bcl-xL levels in patient tumors.

**Fig. 4.**
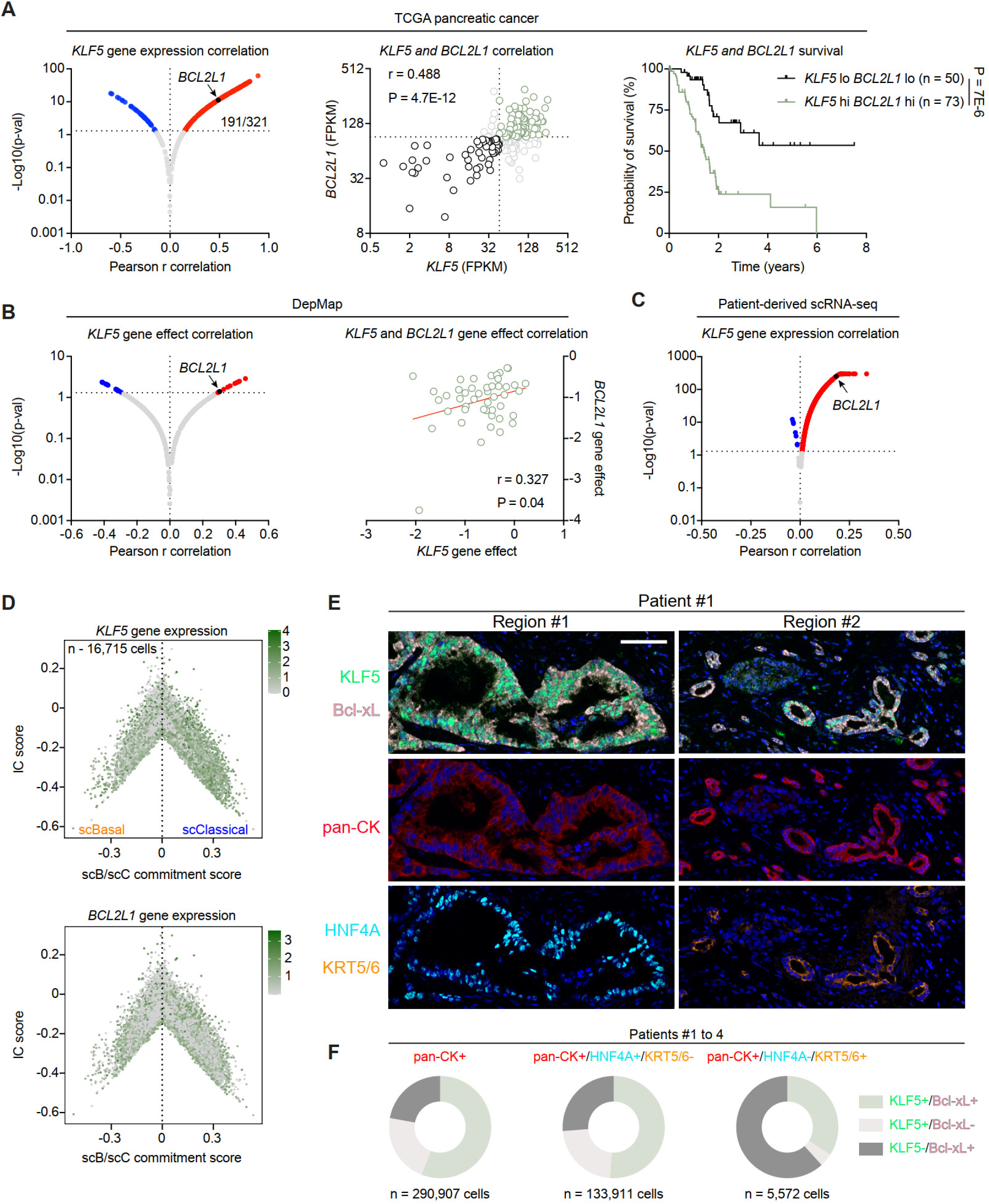
KLF5 and Bcl-xL are co-expressed in patients independent of subtype-identity. (**A**) Pearson r correlation (left) in pancreatic cancer patients from The Cancer Genome Atlas (TCGA) database of *KLF5* gene expression against KLF5-dependent genes in all four cell lines (AsPC1, HPAFII, BxPC3, L3.6pl). *KLF5* and *BCL2L1* gene expression are correlated in patients (middle) and Kaplan-Meier survival analysis of *KLF5*-hi, *BLC2L1*-hi vs *KLF5*-lo, *BLC2L1*-lo patients (right). FPKM cut off of 46.46 and 91.25 were used for *KLF5* and *BCL2L1*, respectively. (**B**) Pearson r correlation (left) of CRISPR-dependency data from DepMap data of KLF5 against KLF5-dependent genes in all four cell lines as described in (A). Correlation of *KLF5* and *BCL2L1* gene effect (right). (**C**) Pearson r correlation of *KLF5* gene expression against KLF5-dependent genes in all four cell lines as described in (A) in patient single-cell RNA-seq (scRNA-seq). (**D**) Cell state diagram on the tumorigenic cells showing *KLF5* and *BCL2L1* expression in scRNA-seq. scBasal-scClassical commitment score (x-axis) and intermediate co-expressor score (y-axis). (**E**) Representative multiplex immunofluorescence image validating KLF5 and Bcl-xL co-expression in patient samples independent of subtype-identity. Scale bar represents 50 µm. (**F**) Quantification of patients 1 to 4 of KLF5 and Bcl-xL co-expression in total tumorigenic cells (pan-CK+), classical cells (pan-CK+, HNF4A+, KRT5/6-) or basal-like cells (pan-CK+, HNF4A-, KRT5/6+).

### Loss of KLF5 induces apoptosis

Decreased cell viability may be due to induction of programmed cell death such as apoptosis, disruption of cell cycle progression or a combination of both. Because we observed decreased Bcl-xL protein levels upon KLF5 knockdown, we hypothesized that depletion of KLF5 would lead to increased apoptosis. In line with this hypothesis, knockdown of KLF5 robustly induced apoptosis in HPAFII, BxPC3 and L3.6pl cells, and modestly in AsPC1 cells, consistent with the effects on cell viability (Fig. 5 A and B). Apoptosis was assessed by a caspase-substrate that upon cleavage by active caspase 3/7 intercalates into DNA and becomes fluorescent. We next sought to understand if apoptosis inhibition rescues KLF5-dependent cell viability. To this end, we depleted KLF5 and treated cells with pan-caspase inhibitor Q-VD-OPh in HPAFII, BxPC3 and L3.6pl cells. Because we did not observe a robust induction of apoptosis in AsPC1 cells, we did not use this cell line in the subsequent studies. We additionally used a live cell annexin V stain to investigate apoptosis inhibition as Q-VD-OPh inhibits caspase activity. Q-VD-OPh treatment resulted in a partial rescue of cell viability across all three cell lines with complete inhibition of apoptosis assessed by annexin V staining (Fig. 5 C to E and fig. S11 A to D). Thus, we propose KLF5 elicits pleiotropic effects on cell viability through additional mechanisms such as cell cycle disruption, consistent with previous reports (*23*). Because pan-caspase inhibition partially rescued cell viability upon KLF5 knockdown, we next assessed whether apoptosis induction can be blocked by Bcl-xL overexpression. Thus, we transiently overexpressed mCherry-tagged Bcl-xL and observed that apoptosis induction following KLF5 depletion could be partially blocked by Bcl-xL overexpression (Fig. 5 F to G and fig. S11 F and G). Because we showed that repression of the downstream KLF5-bound enhancer of *BCL2L1* decreases gene expression, we asked if simultaneously targeting all three enhancers with Zim3-dCas9-mediated repression induces apoptosis and decreases cell viability. We observed an increase in caspase 3/7 activity (Fig. 5 I to K), albeit lower than complete KLF5 depletion (Fig. 5A), and a decrease in cell viability (Fig. 5I and fig. S11E). Lastly, we rapidly degraded KLF5 and treated cells with standard of care chemotherapy gemcitabine/paclitaxel or FOLFIRINOX (5-fluorouracil, irinotecan, oxaliplatin). Interestingly, we found KLF5 degradation sensitized cells to gemcitabine/paclitaxel treatment (Fig. 5 L and M), while only eliciting an additive effect in combination with FOLFIRINOX (fig. S11 H and I). Sensitization to gemcitabine/paclitaxel combination therapy is consistent with previous reports showing Bcl-xL-mediates resistance to these therapies (*24, 25*). Due to the strong effect on cell viability upon KLF5 degradation in BxPC3 and AsPC1 cells (fig. S9 D and E), we were unable to perform combination treatment with gemcitabine/paclitaxel and FOLFIRINOX. Thus, we posit that KLF5 binds multiple enhancers to regulate target genes such as the anti-apoptotic protein *BCL2L1*, which partially contributes to KLF5-mediated induction of apoptosis, to regulate cell viability in a subtype-independent manner in pancreatic cancer.

**Fig. 5.**
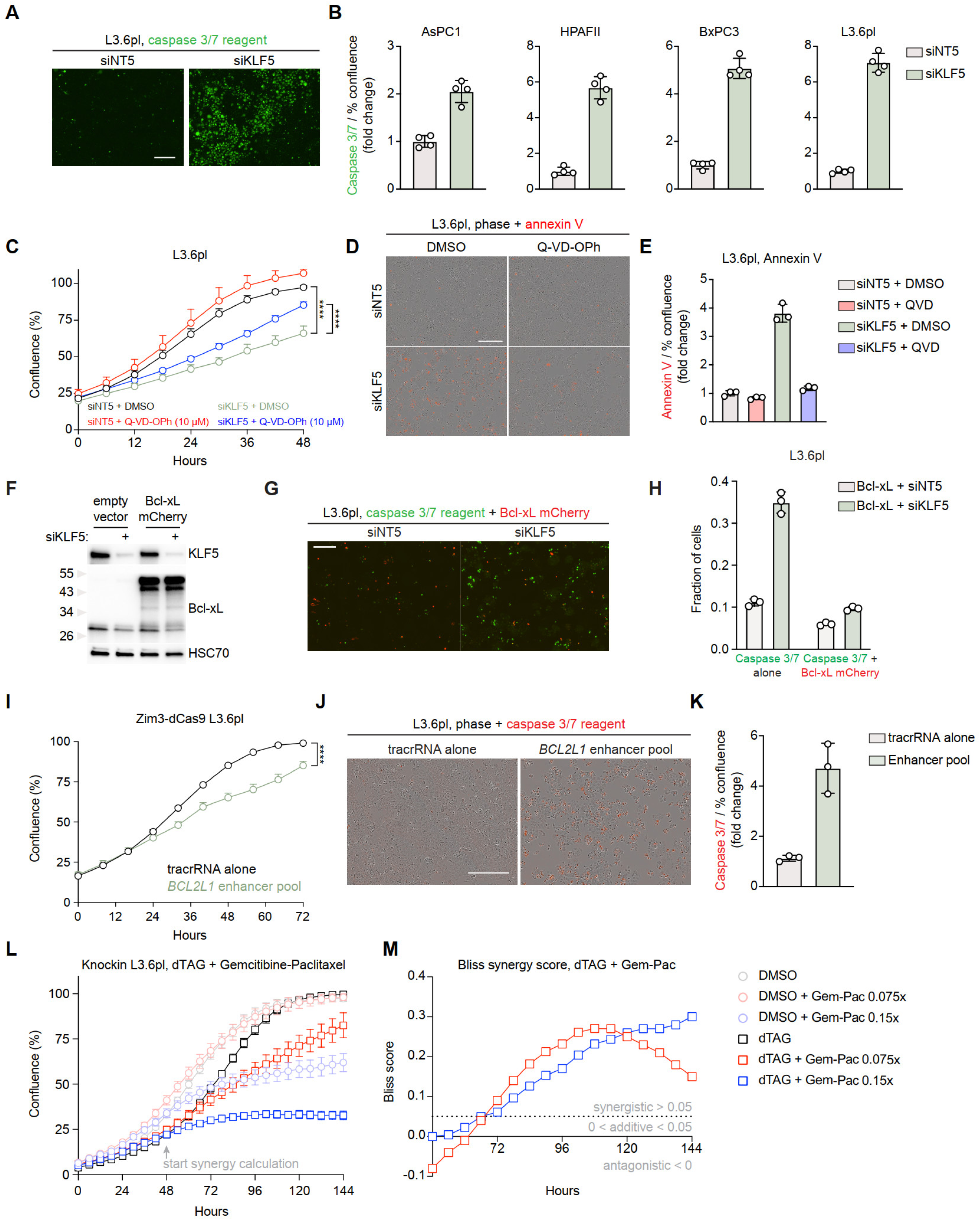
KLF5 regulates apoptosis in pancreatic cancer. (**A and B**) Cells were treated with KLF5 or non-targeting siRNA mix, incubated for 4 h and media was changed with live cell caspase 3/7 reagent (1:1,000) and monitored via live cell imaging for proliferation and green calibrated units (GCU). n = 4 biological replicates. (A) Representative caspase 3/7 induction images, scale bar represents 100 µm. (B) Fold change (siKLF5 vs siNT5) of GCU per % confluency for AsPC1, HPAFII, BxPC3 and L3.6pl cells. (**C to E**) L3.6pl cells were treated with KLF5 or non-targeting siRNA mix, incubated for 4 h and media was changed with DMSO or 10 µM Q-VD-OPh with live cell annexin V reagent (1:1,500) and monitored via live cell imaging for proliferation and red calibrated units (RCU). n = 3 biological replicates. (C) Quantification of confluency over time. Unpaired Student’s t-test on the area under the curve (AUC), **** p < 0.0001. (D) Representative phase and annexin V induction images. Scale bar represents 100 µm. (E) Fold change (siKLF5 vs siNT5) of RCU per % confluency. (**F to H**) L3.6pl cells were transfected with either empty vector backbone (pcDNA3.1) or containing Bcl-xL-mCherry for 24 h and subsequently transfected with KLF5 or non-targeting siRNA mix, incubated for 4 h and media was changed with live cell caspase 3/7 reagent (1:1,000) and monitored via live cell imaging for proliferation and GCU and RCU. n = 3 biological replicates. (F) Representative Western blot analysis of KLF5 and Bcl-xL confirming knockdown and overexpression, respectively. HSC70 was used as a loading control. (G) Representative caspase 3/7 induction and Bcl-xL-mCherry overexpression images. Scale bar represents 100 µm. (H) Fraction of cells with only caspase3/7 reagent (GCU) or caspase 3/7 reagent and mCherry (GCU + RCU). (**I to K**) Lentiviral Zim3-dCas9 L3.6pl cells were treated with a pool targeting downstream enhancers of *BCL2L1* or tracrRNA alone, incubated for 4 h and media was changes with live cell caspase 3/7 reagent (1:1,000) and monitored via live cell imaging for proliferation and RCU. n = 3 biological replicates. (F) Quantification of confluency over time. Unpaired Student’s t-test on the AUC, **** p < 0.0001. (G) Representative caspase 3/7 induction images, scale bare represents 100 µm. (H) Fold change (enhancer pool vs tracrRNA alone) of RCU per % confluency. (**L and M**) Knockin L3.6pl cells were plated overnight and treated with indicated reagents and monitored for confluency. n = 6 biological replicates. (L) Quantification of confluency over time. (M) Quantification of Bliss synergy score from (L).

## Discussion

Through an unbiased bioinformatic analysis of subtype-independent mechanisms of pancreatic cancer cell survival, we identified KLF5 as a pancreatic cancer subtype-independent essential gene. Although KLF5 activation has been implicated in many cancers, the potential utilities of targeting KLF5 or its downstream effectors remains unclear. Through the rapid degradation of KLF5, we identified early enhancers such as those controlling the expression of the anti-apoptotic gene *BCL2L1*. Subsequent enhancer inactivation with Zim3-dCas9 confirmed KLF5-bound enhancers are necessary for target gene expression. Lastly, we confirmed co-expression of KLF5 and Bcl-xL in patient samples independent of subtype identity. Lastly, transcriptomic and epigenetic studies with the non-cancer human pancreatic ductal epithelial cells will help shed light on the role of KLF5-related transcriptional mechanisms in a non-oncogenic context i.e. if their enhancer landscape downstream of the *BCL2L1* locus is active and producing eRNA.

Interestingly, we found that genes rapidly downregulated following KLF5 degradation were connected with multiple KLF5 bound sites displaying higher levels of eRNA transcription compared to either nonregulated genes or late-responsive genes, suggesting KLF5 regulates its target genes through highly transcribed interconnected hubs. These hubs have been shown to be important in tumor biology for resistance to therapy (*13*) and promote aggressiveness in many cancers including acute leukemia (*26*), glioblastoma (*12*) and Ewing sarcoma (*15*). Interestingly, these enhancer elements include, but are not limited to, super enhancers. Multi-connected hubs have been associated with high transcriptional activity, often regulating genes critical for cell identity (*27*), also observed in pancreatic cancer (*28*). Interestingly, these findings suggest that KLF5 plays a central role in controlling the transcription of a select set of important cancer-related genes with their enhancers displaying particularly high levels of transcriptional output.

Our study sheds light on KLF5 as a potential therapeutic in pancreatic cancer. Since both subtypes can exist within the same tumors, future development of Proteolysis Targeting Chimera (PROTAC) targeting KLF5 could unravel new options for therapy. Alternatively, targeting downstream early targets of KLF5 such as Bcl-xL with small molecule inhibitors such as A-1331852 may provide a valuable therapeutic option in *KLF5*-hi/*BCL2L1*-hi tumors. KLF5 recognizes a CG-rich DNA motif and was reported to have different binding sites between different cancers. However, we observed consistent binding between pancreatic cancer cell lines of different subtypes. This suggests small molecule approaches in pancreatic cancer will inhibit KLF5 activity or downstream targets regardless of subtype identity. eRNAs have been shown to facilitate transcription factor recruitment (*29*), mediate Pol II pause release (*30*) and be used as a readout for gene expression (*31*). Another potential therapeutic application of our findings could be the use of antisense oligonucleotides (ASO) targeting the eRNAs of *BCL2L1* to elucidate its role either as a readout for *BCL2L1* expression or influencing KLF5 binding or transcription machinery will be important to modulate KLF5 activity to alter the KLF5-Bcl-xL axis. Together, our findings uncover important new, subtype-independent transcriptional regulatory mechanisms essential for PDAC cell survival. Additional studies targeting these mechanisms *in vivo* will be important for establishing their potential clinical utility.

## Materials and Methods

### Cell lines, cell culture, siRNAs/sgRNAs, plasmids, siRNA/sgRNA transfections, drugs, live cell imaging

AsPC1 and BxPC3 cells were maintained in RPMI 1640 media supplemented with 10% FBS. HPAFII and L3.6pl cells were maintained in phenol-free MEM media supplemented with 10% FBS. HEK293T cells were maintained in DMEM media supplemented with 10% FBS. All cells were cultured with 1% penicillin/streptomycin. Cell cultures were incubated in a humidified 5% CO2 incubator at 37°C. All siRNAs were purchased from Horizon Discovery and all sgRNAs were purchased from GenScript and sequences are listed in Supplementary Table 1. Plasmids were obtained as follows: pMD2.G (Addgene, 12259), psPAX2 (Addgene, 12260), lentiCas9-BLAST (Addgene, 52962), pHR-UCOE-EF1a-Zim3-dCas9-P2A-GFP (Addgene, 188778). pcDNA3.1 empty vector and pcDNA3.1 Bcl-xL mCherry were gifts from Frank Essmann. For siRNA/sgRNA transfections, 7.5 x 10^5^ cells were plated in 1.6 mL of antibiotic-free media in a 6 well plate and 0.4 mL of transfection mix (400 µL OptiMEM (ThermoFisher Scientific, 31985070), 4 µL RNAiMAX (ThermoFisher Scientific, 13778150), 1.5 µL of 20 µM siRNA or sgRNA) was added to the cells for a total of 2 mL media. The efficiency of siRNA knockdown was determined by Western blotting. Drugs were purchased as follows: dTAGV-1 TFA (MedChemExpress, HY-145514), Q-VD-OPh (MedChemExpress, HY-12305), paclitaxel (Selleckchem, S1150), gemcitabine (Selleckchem, S1714), 5-fluorouracil (Selleckchem, S1209), irinotecan (Selleckchem, S1198), oxaliplatin (Selleckchem, S1224). Gemcitabine-paclitaxel 1X mix was made as follows: 130 nM gemcitabine, 14 nM paclitaxel. Folfirinox 1X mix was made as follows: 32 nM 5-fluorouracil, 400 nM irinotecan, 320 nM oxaliplatin. Live cell detection reagents were purchased as follows: Caspase 3/7 (ThermoFisher Scientific, C10423 & C10423), annexin V (Sartorius, 4641). For live cell imaging, cells were plated at 5 x 10^4^ or 2 x 10^4^ in a 24 or 48 well plate, respectively, 24 hr before treatment. Media was changed with indicated drugs and live cell detection reagents (Caspase 3/7, 1:750) or annexin V (1:1500), cultured for 48 hr in the continued presence of the drug and reagents and proliferation and green or red calibrated units (GCU, RCU) were monitored using an Incucyte (Sartorius) by imaging 9 regions of each well every 6-8 hr. Crystal violet stained cells were quantified using Incucyte software. To calculate the Bliss synergy score, first we calculated the Effect (E): 1 – (treated_% confluency_ / control_% confluency_). Then, we used the following formula: E_dTAG + Chemo_ – (E_dTAG_ + E_Chemo_ – (E_dTAG_ x E_Chemo_)). The script for the quantification of green, red and mixed cells (Fig. 5 G and H) can be found here: https://github.com/bhc-rbct/cell-marker-spot-counter.git.

### Knockin cell line generation

For knock-in cell line generation, we performed two transfections as follows. First, 1 x 10^6^ cells were plated in a 6 well plate 24 hr before transfection. 2 μg of lentiCas9-BLAST and 500 ng of construct cloned in a pUC57-mini backbone were forward transfected with Lipofectamine 3000 Transfection Reagent (ThermoFisher Scientific, L3000015) following supplier’s recommendations. Custom constructs were synthesized and cloned into a pUC57-mini backbone from GenScript. Received lyophilized plasmids were resuspended in TE buffer and prepped (Macherey-Nagel, 740412.50). 24 hr post forward transfection, reverse transfection with two sgRNAs targeting the start codon were carried out as described above. Cells were incubated for 48 hr and subsequently transferred to a 10 cm dish and cultured until 70% confluency. The bulk population was generated via fluorescence-activated cell sorting (FACS) with a Sony SH800S cell sorter. Single cell clones were generated by diluting cells to 0.5 cells per 200 µL media and added to a 96 well plate and subsequently expanded.

### Western blot

Cell lysates were prepared by lysing cells in ice-cold RIPA buffer (Serva, 39244). Briefly, cell culture media was removed from adherent cells, washed twice with ice-cold PBS, added RIPA buffer to cells with freshly added 0.1 mM PMSF protease inhibitor (ThermoFisher Scientific, 36978) and 1% v/v phosphatase inhibitor cocktail 3 (Sigma-Aldrich, P0044), incubated for 5 min on a rocker, scraped and transferred to a new tube. Lysates were sonicated for 4 cycles of 30 sec on/off using a Bioruptor Pico (Diagenode). 4x Laemmli Sample Buffer (Bio-Rad, 1610747) with 10% beta-mercaptoethanol was added to samples and ran 20 µL on a 4-20% Mini-PROTEAN TGX Precast Protein Gel (Bio-Rad, 4561095) and transferred to a nitrocellulose membrane (Bio-Rad, 1704158). Membranes were blocked in 5% milk in TBST and incubated with primary antibodies (Supplementary Table 2) overnight at 4°C. Rabbit (Cell Signaling Technology, 7074) and mouse (Cell Signaling Technology, 7076) horseradish peroxidase (HRP)-conjugated secondary antibodies were used. Western blot membranes were developed with Clarity Max Western ECL Substrate (Bio-Rad, 1705062) and chemiluminescence was detected using a ChemiDoc MP Imaging System (Bio-Rad).

### DNA extraction and genotyping PCR

Cell culture medium was removed, washed twice with PBS, lysed with lysis buffer (200 mM NaCl, 5 mM EDTA, 100 mM Tris-HCl, 0.2% v/v SDS), incubated for 5 min on rocker, scraped and transferred to a new tube. Samples were treated with proteinase K (ThermoFisher Scientific, EO0491) and incubated on a ThermoMixer (Eppendorf) at 56°C overnight. Equal volumes of isopropanol were added, vortexed, incubated at −80°C for 1 hr, centrifuged, washed twice with 70% ethanol and resuspended pellet with nuclease free water. PCR was carried out from 200 ng extracted DNA per manufacturer’s protocol with GoTaq DNA Polymerase (Promega, M3001) and PCR nucleotide mix (Promega, C1141).

### Lentiviral production and infection

HEK293T cells were plated 24 hr before transfection to be 90-95% confluent. For a 10 cm plate, 0.72 pmol pMD2, 1.3 pmol psPAX2 and 1.64 pmol plasmid of interest were co-transfected with Lipofectamine 3000 Transfection Reagent following manufacturer’s recommendations. Briefly, media was removed, added transfection mix, incubated for 2 hr and 6 mL of media was added to cells. Supernatant was collected 24, 48 and 72 hr post transfection and pooled. Lentivirus was isolated via centrifugation and supernatant was transferred to a new tube and subsequently concentrated with Lenti-X Concentrator (Takara, 631232) following supplier’s protocol. 5 x 10^5^ cells in a 6 well plate were infected by adding 8 µg/mL final concentration polybrene (Santa Cruz Biotechnology, sc-134220), 100 µL of concentrated virus, and centrifuging for 1000 g for 2 hr at 30°C. Infected cells were sorted for GFP (Zim3-dCas9 infected cells) and expanded.

### ChIP and ChIP-seq library preparation

Chromatin immunoprecipitation was performed by crosslinking cells with 1% formaldehyde for 20 min and quenched by 1.25 M glycine for 5 min. Cells were lysed with lysis buffer (150 mM NaCl, 20 mM EDTA, 50 mM Tris-HCl, 0.5% v/v NP-40, 1% v/v Triton X-100, 20 mM NaF). Nuclear pellets were resuspended and sonicated in sonication buffer (150 mM NaCl, 20 mM EDTA, 50 mM Tris-HCl, 1% v/v NP-40, 0.5% v/v sodium deoxycholate, 20 mM NaF, 0.1% SDS) using a Bioruptor Pico (Diagenode) and a cycle setting of 30 sec on/off until fragment sizes around 200-500 bp was reached. Protease (Roche, 11836153001) inhibitors were added to both lysis and sonication buffers. Samples were precleared by 50% slurry of sepharose (Cytiva, 17012001) resuspended in sonication buffer. Antibodies were added and incubated rotating overnight (Supplementary Table 2). Sepharose beads coupled with Protein A (Cytiva, 17078001, rabbit antibodies) or Protein G (Cytiva, 17061801, mouse or goat antibodies) were added to samples and incubated for 2 hr, washed, de-crosslinked, RNAse A (ThermoFisher Scientific, EN0531) and proteinase K treated, and DNA was extracted. For qPCR validation, 10 µL of extracted DNA was diluted 1:4 in nuclease free water and 2 µL was used in the final reaction. ChIP DNA was quantified with Qubit (Invitrogen) using the Qubit 1X dsDNA High Sensitivity Kit (ThermoFisher Scientific, Q33231) and libraries were prepped per supplier’s protocol using the MicroPlex Library Preparation Kit v3 (Diagenode, C05010001). The library fragment size was determined by TapeStation (Agilent) using the High Sensitivity D1000 Sample Buffer (Agilent, 5190-6504) and High Sensitivity D1000 ScreenTape (Agilent, 5067-5584). Libraries were sequenced paired end at the Sequencing Core at Robert Bosch Center for Tumor Diseases (Illumina, NextSeq 2000).

### ChIP-seq bioinformatic analysis

Paired end sequencing reads were mapped to the reference genome assembly hg38 using bowtie2 v2.5.2 (*32*). chrM, chrUn, alt and random chromosomes were removed from bam files with ‘grep -v’ command. PCR duplicates were removed with samtools markdup (samtools v1.21) (*33*) and replicates were merged using samtools merge. Bigwig files for individual and merged bam files were generated ignoring duplicates and blacklist regions (*34*) using bamCoverage (deeptools v3.5.5) (*35*) using reads per genomic content (RPGC) normalization. Localization profiles were viewed using the Integrated Genomics Viewer (IGV v2.19.4) (*36*). MACS3 callpeak (macs3 v3.0.1) (*37*) was used to call the significant peaks with --broad cutoff 0.05 for histone marks and non-broad cutoff of 0.05 for transcription factors while using input files from respective conditions as background. ChIP occupancy was evaluated by computeMatrix (deeptools v3.5.5) and the average profiles and heatmaps were generated based on computeMatrix values with plotHeatmap or plotProfile (deeptools v3.5.5). To generate Log2FC bigwigs, we used bigwigCompare (deeptools v3.5.5). hg38 TSS and TES coordinates were retrieved from UCSC table browser. To ascertain TSS specific regions in our system, we intersected TSS coordinates with our H3K4me3 bed file (macs3 callpeak output) to retrieve active TSS using bedtools v2.31.1 (*38*). Differential binding analysis was performed using DiffBind (*39*) (R v4.5.0). Peak annotation was performed using ChIPseeker (*40*) (R v4.5.0).

### RNA extraction, cDNA synthesis, and qPCR

Cells were washed twice with PBS after removing cell culture medium and total RNA was extracted from cells by adding 700 µL of QIAzol, incubated for 5 min on rocker, scraped and transferred to a new tube. 140 µL chloroform was added to the samples, vortexed for 15 sec, incubated for 3 min at room temperature and centrifuged. Aqueous phase was transferred to a new tube and mixed with an equal volume of isopropanol. Samples were incubated at −80°C for 1 hr and centrifuged. RNA pellet was washed twice with 70% ethanol and the pellet was air-dried at room temperature for 5-10 min. 50 µL of nuclease free water was used to resuspend the pellet and concentration was measured using the Nanodrop (DeNovix DS-11). cDNA was synthesized from 500 ng of total RNA using PrimeScript RT Reagent Kit (Takara, RR037A) with a final reaction volume of 10 µL containing oligo(dT) and random hexamers. cDNA was diluted to 50 µL with nuclease free water. For cDNA synthesis of eRNAs, samples were first treated with DNAse I (ThermoFisher Scientific, EN0521) prior to cDNA synthesis and only random hexamers were used. qPCR was performed in duplicate for each sample using 2 µL template in a final volume of 10 µL on a CFX Duet Real-Time PCR System (Bio-Rad) using SsoAdvanced Universal SYBR Green Supermix (Bio-Rad, 1725274). *ACTB* was used to normalize mRNA and eRNA expression. All primers were purchased from Integrated DNA Technologies (IDT) and are listed in Supplementary Table 1.

### RNA-seq bioinformatic analysis

Before RNA-sequencing, RNA integrity was validated by gel electrophoresis and libraries were prepped using 500 ng following supplier’s protocol – Illumina Stranded mRNA Prep, Ligation (Illumina, 20040534). Library concentration and size was determined as described above. Libraries were sent to the Sequencing Core at Robert Bosch Center for Tumor Diseases (Illumina, NextSeq 2000) and sequenced stranded paired end. Paired end sequencing reads were mapped to the reference genome assembly hg38 using STAR (star v2.7.9a) (*41*). Reverse (sense strand) reads were quantified using htseq-count (htseq v2.0.5) (*42*) and used for differential gene expression analysis via DESeq2 (*43*) (R v4.5.0). Gene set enrichment analysis (GSEA v4.2.2) (*44*) was performed with default settings using normalized counts from DESeq2 for expressed genes. Heatmaps were generated with pHeatmap (R v4.5.0) on vst transformed normalized counts from DESeq2. PCA plots were generated with varianceStabilizingTransformation (R v4.5.0) and plotPCA (R v4.5.0).

### ATAC-seq

For ATAC-Seq library preparation, 75,000 cells were harvested and libraries were produced using the ATAC-seq kit (Active Motif, 53150) following manufacturer’s instructions. Library concentration and size was determined as described above. Libraries were sequenced paired end at Sequencing Core at Robert Bosch Center for Tumor Diseases (Illumina, NextSeq 2000).

### ATAC-seq bioinformatic analysis

Paired end sequencing reads were filtered for adapter contamination with trim galore v0.6.1 (https://github.com/FelixKrueger/TrimGalore.git) and mapped to the reference genome assembly hg38 using bowtie2 v2.5.2 with --dovetail. chrM, chrUn, alt and random chromosomes were removed from bam files with ‘grep -v’ command and BamTools Filter (*45*) was used for removing low quality (mapQ ≥ 30). Resulting bam file was converted to a bed file with bedtools v2.31.1 bamtobed. Peak calling and bedgraph generation were performed using macs3 callpeak (macs3 v3.0.1) using shift and extend method to center peaks at 5’ cut site (--nomodel --shift -100 --extsize 200). The data was visualized by using bedGraphToBigWig. To normalize the data, we generated reads in peaks “RiP” with DiffBind (method = DBA_DESEQ2, normalize = DBA_NORM_LIB, library = DBA_LIBSIZE_PEAKREADS, background = FALSE). Normalization factors were generated by 1/(RiP value).

### Click-iT nascent RNA capture and library preparation

Nascent RNA capture was performed by plating 5 x 10^5^ cells in a 6 well plate, incubated overnight and treated with dTAGV-1 for 0, 1, 4 or 24 hr. 0.5 mM 5-ethynyl uridine (EU) in 2 mL of cell culture media was added to cells during the last 1 hr of dTAGV-1 treatment. RNA extraction was performed as described above and 2 µg of isolated RNA was used for the click reaction using the Click-iT Nascent RNA Capture Kit (ThermoFisher Scientific, C10365) following supplier’s protocol. Following the click reaction, RNA was extracted and 500 ng of isolated RNA was used for binding to Dynabeads MyOne Streptavidin T1 beads, washed and proceeded immediately to library preparation using the Universal Plus Total RNA-Seq with NuQuant kit (Tecan Life Sciences, M01523). Library concentration and size was determined as described above. Libraries were sequenced paired end at Sequencing Core at Robert Bosch Center for Tumor Diseases (Illumina, NextSeq 2000).

### Click-iT nascent RNA capture bioinformatic analysis

Paired end sequencing reads were mapped to the reference genome assembly hg38 using bowtie2 v2.5.2. chrM, chrUn, alt and random chromosomes were removed from bam files with ‘grep -v’ command. PCR duplicates were removed with samtools markdup (samtools v1.21) and replicates were merged using samtools merge. Forward and reverse bigwig files for individual and merged bam files were generated ignoring duplicates and blacklist regions (*34*) using bamCoverage (deeptools v3.5.5) using counts per million (cpm) normalization and --filterRNAstrand forward and reverse. Localization profiles were viewed using the Integrated Genomics Viewer (IGV v2.19.4). Occupancy was evaluated by computeMatrix (deeptools v3.5.5) and the average profiles and heatmaps were generated based on computeMatrix values with plotProfile (deeptools v3.5.5).

### H3K4me3 HiChIP bioinformatic analysis

HiChIP was analyzed as previously described (*16*). Briefly, sequencing reads were mapped with HiC-Pro v3.1.0 (*46*) and FitHiChIP v11.0 (*47*) was utilized to generate bedpe file format. hicpro2juicebox (HiC-Pro v3.1.0) was used to generate a ‘.hic’ matrix file which was uploaded to the Tinker platform at Axiotl (https://docs.axiotl.com/) to generate contact maps (Fig. 2H). Counts per million were used to normalize samples. We generated an in-house script (https://github.com/bhc-rbct/Filter-A2A-loops-for-active-genes) which extracts the active TSS of target genes using a cell line specific H3K4me3 bed file (anchor 1) which interacts with (enhancer) regions of interest (anchor 2) i.e. KLF5 regions that interact with KLF5-dependent genes.

### PRO-seq and leChRO-seq bioinformatic analysis

PRO-seq and leChRO-seq were analyzed as previously described (*16*). Briefly, paired end sequencing reads were mapped to hg38 genome using proseq2.0 pipeline (*48*). For PRO-seq, we used the following parameters (-PE --RNA5=R1_5prime --map5=TRUE --opposite-strand=TRUE -- UMI1=6 --UMI2=6) and used the following parameters for leChRO-seq (-SE -G --UMI1=6). Discriminative regulatory element detection (dREG) was generated via https://dreg.dnasequence.org from the bigwigs generated from proseq2.0 pipeline.

### Identification of subtype-independent highly interactive enhancers

To generate distal open, active regions devoid of CTCF binding in AsPC1 and L3.6pl cells, we first extended ATAC summits to 200 bp (+/- 100 bp, referred to as ATAC peaks). Next, we extended annotated TSS coordinates to 8 kb (+/- 4 kb). We chose 8 kb in total as this results in 2 HiChIP bins on either side of the TSS. We then subtracted TSS regions from ATAC peaks to garner distal open regions. These distal open regions were then intersected with H3K27ac peaks to obtain distal open, active regions. We then extended CTCF summits to 200 bp (+/- 100 bp, referred to as CTCF peaks). We then subtracted distal open, active regions with CTCF peaks to obtain distal open, active regions devoid of CTCF binding. Lastly, we intersected the resulting bed files between AsPC1 and L3.6pl to gain subtype-independent distal open, active regions.

To find highly interactive enhancers in both cell systems, we utilized an in-house script (https://github.com/bhc-rbct/Filter-A2A-loops-for-active-genes.git) which extracts the active TSS of target genes using a cell line specific H3K4me3 bed file (anchor 1) which interacts with regions of interest (anchor 2) i.e. all H3K4me3-marked genes to subtype-independent distal open, active regions. Resulting bedpe files from AsPC1 and L3.6pl were intersected to compare identical enhancer-promoter pairs. Enhancer-promoter pairs with contact counts ≥5 were kept and the enhancers were subjected to motif analysis with findMotifsGenome (HOMER v4.11) using default settings (*49*).

### Multispectral imaging

A seven-color multiplex immunofluorescence staining was performed using Opal manual detection kit (Akoya Biosciences, NEL861001KT) following manufacturer’s protocol. The formalin-fixed, paraffin-embedded (FFPE) tumor samples were cut into 5 µm sections and were further deparaffinized, rehydrated, subjected to heat-induced epitope retrieval and incubated with primary antibodies. See Supplementary Table 2 for antibodies. Antibodies are listed in order they were added to the sample. Antibodies were visualized with the following tyramide dyes from the Opal Detection Kit: Opal Polaris 480, Opal 520, Opal 570, Opal 620, Opal 690, and DIG-Opal 780. Slides were mounted with ProLong Diamond Antifade Mountant (ThermoFisher Scientific, P36965) and further imaged using a PhenoImager Fusion system (Akoya Biosciences). Quantification of staining was performed with QuPath v0.5.1 and the script can be found here: https://github.com/bhc-rbct/fluorescence_image_cell_marker_quantification.git. Briefly, we log transformed the mean intensity of each marker per cell and utilized a Gaussian mixture model (GMM) to calculate the minimum intensity to be considered for analysis across all samples.

### Public scRNA-seq analysis

For the scRNA-seq analysis, the following publicly available PDAC patient datasets were downloaded from the GEO database: GSE154778 (*20*), GSE111672 (*18*), GSE155698 (*17*), GSE141017 (*22*), 10.5281/zenodo.3969339 (*19*) and 10.5281/zenodo.6024273 (*21*). These datasets were processed to combine and prepare them for downstream analysis by following the same bioinformatic methodology described in Chijimatsu et al. In brief, Seurat (*50*) objects were created for individual datasets by reading the Cellranger output files into the R environment using the Read10x function and transformed using the CreateSeuratObject function (R v4.2.3 and Seurat v4.3.0.1). Counts of transcripts measured as UMIs were normalized to 10,000 counts per cell and log transformed. Cells with high mitochondrial genes (>25%) were filtered in the QC steps. Other QC metrics for individual datasets regarding UMI and expressed genes were followed same as mentioned in Chijimatsu et al. Datasets were batch-corrected and data integration was performed using the rPCA method according to the Seurat package. Each dataset was scaled and the FindVariableFeatures function was used to find highly variable genes. These genes were used to perform PCA analysis. An anchor was created using FindIntegrationAnchors with arguments that 30 principal components, rPCA, two reference datasets (PRJCA001063 and GSE155698), followed by six datasets, were integrated using the IntegrateData. The integrated dataset was scaled, PCA analysis was done and UMAP was visualized. Cell-type annotation was transferred from the reference dataset BioProject:PRJCA001063.

### Cell State Diagram Analysis

The R code for the cell state diagram was kindly provided by Dr. Peter Winter from Prof. Alex Shalek’s lab at MIT; also described in Raghavan et al. (*3*). Briefly, only ductal cell type II (tumor cells) and molecular subtype classification from Chan-Seng-Yue et al. (*51*) were used in the analysis. Basal gene scores were first transformed into negative values and termed as flipped basal. This negative basal score was added to classical score values to calculate tumor-type scores. The absolute values of the tumor-type score were subtracted from both high correlation scores to calculate an undifferentiated or intermediate score for each cell. These scores were plotted in a 2D format with the X-axis differentiating between basal or classical cell types. The Y-axis determines the intermediate state of the cells. Individual gene scores were calculated and projected onto the wishbone plot.

### Ethics approval

Samples for multiplex immunofluorescence were collected by the Center for Cell Signaling in Gastroenterology (C-SiG) at Mayo Clinic (IRB 21-004887) and IRBs 66-06, 354-06, 19-012104. Informed consent was obtained before participants enrolled in the study.

### Statistical analysis

All bar graphs are represented as mean ± standard deviation (SD) and live cell imaging over time is represented as mean ± standard error mean (SEM). Statistical analyses were performed using GraphPad Prism 10 and R software. Statistical analyses were performed using two-tailed, unpaired Student’s t test and Mann-Whitney test for comparisons between two groups. For live cell imaging involving multiple groups, area under curve (AUC), SEM and degrees of freedom for each condition were calculated and subsequent one-way analysis of variance (ANOVA) on the AUC with Tukey’s multiple comparisons was used. Significance of Kaplan-Meier survival curves was estimated by log-rank (Mantel-Cox) test. For correlation data, two-tailed Pearson correlation coefficients were calculated.

## Acknowledgements

We thank L Morgan and team at Axiotl for their support with HiChIP data analysis. We thank P Winter for bioinformatic assistance generating scRNA-seq PDAC cell state diagrams. We thank A Amoateng at the Robert Bosch Center for Tumor Diseases sequencing facility. We thank K Willecke and A Kechter for technical assistance with mIF.

## Author contributions

TLE and SAJ conceptualized the project and designed experiments. TLE, BJS, ZD, and YS performed experiments. JT and MD established and performed mIF. MM conducted scRNA-seq analysis. NS analyzed mIF and generated bioinformatic scripts. AMA and MJT provided patient material. TLE analyzed data and performed bioinformatic analysis. TLE and SAJ wrote the manuscript.

## Data and materials availability

All sequencing data generated in this study are available in a public, open access repository Sequence Read Archive (PRJNA1283515) and Gene Expression Omnibus: ChIP-seq (GSE301284), mRNA-seq (GSE301283), ATAC-seq (GSE301272), nascent RNA-seq (GSE301285). Publicly available data used in this study include ATAC-seq (GSE271731), ChIP-seq (GSE271732, GSE115463), PRO-seq (GSE271735), leChRO-seq (GSE271734), HiChIP-seq (GSE271733), scRNA-seq (GSE154778, GSE111672, GSE155698, GSE141017, 10.5281/zenodo.3969339, 10.5281/zenodo.6024273.

## References

1. R. L. Siegel, A. N. Giaquinto, A. Jemal, Cancer statistics, 2024. CA Cancer J Clin 74, 12–49 (2024).

2. G. M. O’Kane, B. T. Grunwald, G. H. Jang, M. Masoomian, S. Picardo, R. C. Grant, R. E. Denroche, A. Zhang, Y. Wang, B. Lam, P. M. Krzyzanowski, I. M. Lungu, J. M. S. Bartlett, M. Peralta, F. Vyas, R. Khokha, J. Biagi, D. Chadwick, S. Ramotar, S. Hutchinson, A. Dodd, J. M. Wilson, F. Notta, G. Zogopoulos, S. Gallinger, J. J. Knox, S. E. Fischer, GATA6 Expression Distinguishes Classical and Basal-like Subtypes in Advanced Pancreatic Cancer. Clin Cancer Res 26, 4901–4910 (2020).

3. S. Raghavan, P. S. Winter, A. W. Navia, H. L. Williams, A. DenAdel, K. E. Lowder, J. Galvez-Reyes, R. L. Kalekar, N. Mulugeta, K. S. Kapner, M. S. Raghavan, A. A. Borah, N. Liu, S. A. Vayrynen, A. D. Costa, R. W. S. Ng, J. Wang, E. K. Hill, D. Y. Ragon, L. K. Brais, A. M. Jaeger, L. F. Spurr, Y. Y. Li, A. D. Cherniack, M. A. Booker, E. F. Cohen, M. Y. Tolstorukov, I. Wakiro, A. Rotem, B. E. Johnson, J. M. McFarland, E. T. Sicinska, T. E. Jacks, R. J. Sullivan, G. I. Shapiro, T. E. Clancy, K. Perez, D. A. Rubinson, K. Ng, J. M. Cleary, L. Crawford, S. R. Manalis, J. A. Nowak, B. M. Wolpin, W. C. Hahn, A. J. Aguirre, A. K. Shalek, Microenvironment drives cell state, plasticity, and drug response in pancreatic cancer. Cell 184, 6119–6137 e6126 (2021).

4. M. Ema, D. Mori, H. Niwa, Y. Hasegawa, Y. Yamanaka, S. Hitoshi, J. Mimura, Y. Kawabe, T. Hosoya, M. Morita, D. Shimosato, K. Uchida, N. Suzuki, J. Yanagisawa, K. Sogawa, J. Rossant, M. Yamamoto, S. Takahashi, Y. Fujii-Kuriyama, Kruppel-like factor 5 is essential for blastocyst development and the normal self-renewal of mouse ESCs. Cell Stem Cell 3, 555–567 (2008).

5. P. He, J. W. Yang, V. W. Yang, A. B. Bialkowska, Kruppel-like Factor 5, Increased in Pancreatic Ductal Adenocarcinoma, Promotes Proliferation, Acinar-to-Ductal Metaplasia, Pancreatic Intraepithelial Neoplasia, and Tumor Growth in Mice. Gastroenterology 154, 1494–1508 e1413 (2018).

6. Y. Liu, B. Guo, E. Aguilera-Jimenez, V. S. Chu, J. Zhou, Z. Wu, J. M. Francis, X. Yang, P. S. Choi, S. D. Bailey, X. Zhang, Chromatin Looping Shapes KLF5-Dependent Transcriptional Programs in Human Epithelial Cancers. Cancer Res 80, 5464–5477 (2020).

7. C. M. Uyehara, E. Apostolou, 3D enhancer-promoter interactions and multi-connected hubs: Organizational principles and functional roles. Cell Rep 42, 112068 (2023).

8. G. R. Diaferia, C. Balestrieri, E. Prosperini, P. Nicoli, P. Spaggiari, A. Zerbi, G. Natoli, Dissection of transcriptional and cis-regulatory control of differentiation in human pancreatic cancer. EMBO J 35, 595–617 (2016).

9. X. Zhang, P. S. Choi, J. M. Francis, G. F. Gao, J. D. Campbell, A. Ramachandran, Y. Mitsuishi, G. Ha, J. Shih, F. Vazquez, A. Tsherniak, A. M. Taylor, J. Zhou, Z. Wu, A. C. Berger, M. Giannakis, W. C. Hahn, A. D. Cherniack, M. Meyerson, Somatic Superenhancer Duplications and Hotspot Mutations Lead to Oncogenic Activation of the KLF5 Transcription Factor. Cancer Discov 8, 108–125 (2018).

10. J. D. Campbell, A. Alexandrov, J. Kim, J. Wala, A. H. Berger, C. S. Pedamallu, S. A. Shukla, G. Guo, A. N. Brooks, B. A. Murray, M. Imielinski, X. Hu, S. Ling, R. Akbani, M. Rosenberg, C. Cibulskis, A. Ramachandran, E. A. Collisson, D. J. Kwiatkowski, M. S. Lawrence, J. N. Weinstein, R. G. Verhaak, C. J. Wu, P. S. Hammerman, A. D. Cherniack, G. Getz, N. Cancer Genome Atlas Research, M. N. Artyomov, R. Schreiber, R. Govindan, M. Meyerson, Distinct patterns of somatic genome alterations in lung adenocarcinomas and squamous cell carcinomas. Nat Genet 48, 607–616 (2016).

11. X. Shen, Y. Zhang, Z. Xu, H. Gao, W. Feng, W. Li, Y. Miao, Z. Xu, Y. Zong, J. Zhao, A. Lu, KLF5 inhibition overcomes oxaliplatin resistance in patient-derived colorectal cancer organoids by restoring apoptotic response. Cell Death Dis 13, 303 (2022).

12. S. L. Breves, D. C. Di Giammartino, J. Nicholson, S. Cirigliano, S. R. Mahmood, U. J. Lee, A. Martinez-Fundichely, J. Jungverdorben, R. Singhania, S. Rajkumar, R. Kirou, L. Studer, E. Khurana, A. Polyzos, H. A. Fine, E. Apostolou, Three-dimensional regulatory hubs support oncogenic programs in glioblastoma. Mol Cell, (2025).

13. F. H. Hamdan, A. M. Abdelrahman, A. P. Kutschat, X. Wang, T. L. Ekstrom, N. Jalan-Sakrikar, C. Wegner Wippel, N. Taheri, L. Tamon, W. Kopp, J. Aggrey-Fynn, A. V. Bhagwate, R. Alva-Ruiz, I. Lynch, J. Yonkus, R. L. Kosinsky, J. Gaedcke, S. A. Hahn, J. T. Siveke, R. Graham, Z. Najafova, E. Hessmann, M. J. Truty, S. A. Johnsen, Interactive enhancer hubs (iHUBs) mediate transcriptional reprogramming and adaptive resistance in pancreatic cancer. Gut, (2023).

14. A. Swaroop, F. Yue, Chromatin hubs drive key regulatory networks in leukemia. Mol Cell 85, 1–2 (2025).

15. R. Sanalkumar, R. Dong, L. Lee, Y. H. Xing, S. Iyer, I. Letovanec, S. La Rosa, G. Finzi, E. Musolino, R. Papait, I. Chebib, G. P. Nielsen, R. Renella, G. M. Cote, E. Choy, M. Aryee, K. Stegmaier, I. Stamenkovic, M. N. Rivera, N. Riggi, Highly connected 3D chromatin networks established by an oncogenic fusion protein shape tumor cell identity. Sci Adv 9, eabo3789 (2023).

16. T. L. Ekstrom, R. M. Rosok, A. M. Abdelrahman, C. Parassiadis, M. Manjunath, M. Y. Dittrich, X. Wang, A. P. Kutschat, A. Kanakan, A. Rajput, N. Schacherer, T. Lukic, D. M. Carlson, J. Thiel, W. Kopp, P. Stroebel, V. Ellenrieder, J. Gaedcke, M. Dong, Z. Najafova, M. J. Truty, E. Hessmann, S. A. Johnsen, Glucocorticoid receptor suppresses GATA6-mediated RNA polymerase II pause release to modulate classical subtype identity in pancreatic cancer. Gut, (2025).

17. N. G. Steele, E. S. Carpenter, S. B. Kemp, V. R. Sirihorachai, S. The, L. Delrosario, J. Lazarus, E. D. Amir, V. Gunchick, C. Espinoza, S. Bell, L. Harris, F. Lima, V. Irizarry-Negron, D. Paglia, J. Macchia, A. K. Y. Chu, H. Schofield, E. J. Wamsteker, R. Kwon, A. Schulman, A. Prabhu, R. Law, A. Sondhi, J. Yu, A. Patel, K. Donahue, H. Nathan, C. Cho, M. A. Anderson, V. Sahai, C. A. Lyssiotis, W. Zou, B. L. Allen, A. Rao, H. C. Crawford, F. Bednar, T. L. Frankel, M. Pasca di Magliano, Multimodal Mapping of the Tumor and Peripheral Blood Immune Landscape in Human Pancreatic Cancer. Nat Cancer 1, 1097–1112 (2020).

18. R. Moncada, D. Barkley, F. Wagner, M. Chiodin, J. C. Devlin, M. Baron, C. H. Hajdu, D. M. Simeone, I. Yanai, Integrating microarray-based spatial transcriptomics and single-cell RNA-seq reveals tissue architecture in pancreatic ductal adenocarcinomas. Nat Biotechnol 38, 333–342 (2020).

19. J. Peng, B. F. Sun, C. Y. Chen, J. Y. Zhou, Y. S. Chen, H. Chen, L. Liu, D. Huang, J. Jiang, G. S. Cui, Y. Yang, W. Wang, D. Guo, M. Dai, J. Guo, T. Zhang, Q. Liao, Y. Liu, Y. L. Zhao, D. L. Han, Y. Zhao, Y. G. Yang, W. Wu, Single-cell RNA-seq highlights intra-tumoral heterogeneity and malignant progression in pancreatic ductal adenocarcinoma. Cell Res 29, 725–738 (2019).

20. W. Lin, P. Noel, E. H. Borazanci, J. Lee, A. Amini, I. W. Han, J. S. Heo, G. S. Jameson, C. Fraser, M. Steinbach, Y. Woo, Y. Fong, D. Cridebring, D. D. Von Hoff, J. O. Park, H. Han, Single-cell transcriptome analysis of tumor and stromal compartments of pancreatic ductal adenocarcinoma primary tumors and metastatic lesions. Genome Med 12, 80 (2020).

21. R. Chijimatsu, S. Kobayashi, Y. Takeda, M. Kitakaze, S. Tatekawa, Y. Arao, M. Nakayama, N. Tachibana, T. Saito, D. Ennishi, S. Tomida, K. Sasaki, D. Yamada, Y. Tomimaru, H. Takahashi, D. Okuzaki, D. Motooka, T. Ohshiro, M. Taniguchi, Y. Suzuki, K. Ogawa, M. Mori, Y. Doki, H. Eguchi, H. Ishii, Establishment of a reference single-cell RNA sequencing dataset for human pancreatic adenocarcinoma. iScience 25, 104659 (2022).

22. Y. Schlesinger, O. Yosefov-Levi, D. Kolodkin-Gal, R. Z. Granit, L. Peters, R. Kalifa, L. Xia, A. Nasereddin, I. Shiff, O. Amran, Y. Nevo, S. Elgavish, K. Atlan, G. Zamir, O. Parnas, Single-cell transcriptomes of pancreatic preinvasive lesions and cancer reveal acinar metaplastic cells’ heterogeneity. Nat Commun 11, 4516 (2020).

23. C. Rogerson, S. Ogden, E. Britton, O. Consortium, Y. Ang, A. D. Sharrocks, Repurposing of KLF5 activates a cell cycle signature during the progression from a precursor state to oesophageal adenocarcinoma. Elife 9, (2020).

24. D. Thummuri, S. Khan, P. W. Underwood, P. Zhang, J. Wiegand, X. Zhang, V. Budamagunta, A. Sobh, A. Tagmount, A. Loguinov, A. N. Riner, A. S. Akki, E. Williamson, R. Hromas, C. D. Vulpe, G. Zheng, J. G. Trevino, D. Zhou, Overcoming Gemcitabine Resistance in Pancreatic Cancer Using the BCL-X(L)-Specific Degrader DT2216. Mol Cancer Ther 21, 184–192 (2022).

25. A. M. Ibrado, L. Liu, K. Bhalla, Bcl-xL overexpression inhibits progression of molecular events leading to paclitaxel-induced apoptosis of human acute myeloid leukemia HL-60 cells. Cancer Res 57, 1109–1115 (1997).

26. G. Gambi, F. Boccalatte, J. Rodriguez Hernaez, Z. Lin, B. Nadorp, A. Polyzos, J. Tan, K. Avrampou, G. Inghirami, A. Kentsis, E. Apostolou, I. Aifantis, A. Tsirigos, 3D chromatin hubs as regulatory units of identity and survival in human acute leukemia. Mol Cell 85, 42–60 e47 (2025).

27. A. M. Oudelaar, J. O. J. Davies, L. L. P. Hanssen, J. M. Telenius, R. Schwessinger, Y. Liu, J. M. Brown, D. J. Downes, A. M. Chiariello, S. Bianco, M. Nicodemi, V. J. Buckle, J. Dekker, D. R. Higgs, J. R. Hughes, Single-allele chromatin interactions identify regulatory hubs in dynamic compartmentalized domains. Nat Genet 50, 1744–1751 (2018).

28. X. Wang, A. P. Kutschat, J. Aggrey-Fynn, F. H. Hamdan, R. P. Graham, A. Q. Wixom, Y. Souto, S. Ladigan-Badura, J. A. Yonkus, A. M. Abdelrahman, R. Alva-Ruiz, J. Gaedcke, P. Strobel, R. L. Kosinsky, F. Wegwitz, P. Hermann, M. J. Truty, J. T. Siveke, S. A. Hahn, E. Hessmann, S. A. Johnsen, Z. Najafova, Identification of a DeltaNp63-Dependent Basal-Like A Subtype-Specific Transcribed Enhancer Program (B-STEP) in Aggressive Pancreatic Ductal Adenocarcinoma. Mol Cancer Res, (2023).

29. A. A. Sigova, B. J. Abraham, X. Ji, B. Molinie, N. M. Hannett, Y. E. Guo, M. Jangi, C. C. Giallourakis, P. A. Sharp, R. A. Young, Transcription factor trapping by RNA in gene regulatory elements. Science 350, 978–981 (2015).

30. V. Gorbovytska, S. K. Kim, F. Kuybu, M. Gotze, D. Um, K. Kang, A. Pittroff, T. Brennecke, L. M. Schneider, A. Leitner, T. K. Kim, C. D. Kuhn, Enhancer RNAs stimulate Pol II pause release by harnessing multivalent interactions to NELF. Nat Commun 13, 2429 (2022).

31. E. Arner, C. O. Daub, K. Vitting-Seerup, R. Andersson, B. Lilje, F. Drablos, A. Lennartsson, M. Ronnerblad, O. Hrydziuszko, M. Vitezic, T. C. Freeman, A. M. Alhendi, P. Arner, R. Axton, J. K. Baillie, A. Beckhouse, B. Bodega, J. Briggs, F. Brombacher, M. Davis, M. Detmar, A. Ehrlund, M. Endoh, A. Eslami, M. Fagiolini, L. Fairbairn, G. J. Faulkner, C. Ferrai, M. E. Fisher, L. Forrester, D. Goldowitz, R. Guler, T. Ha, M. Hara, M. Herlyn, T. Ikawa, C. Kai, H. Kawamoto, L. M. Khachigian, S. P. Klinken, S. Kojima, H. Koseki, S. Klein, N. Mejhert, K. Miyaguchi, Y. Mizuno, M. Morimoto, K. J. Morris, C. Mummery, Y. Nakachi, S. Ogishima, M. Okada-Hatakeyama, Y. Okazaki, V. Orlando, D. Ovchinnikov, R. Passier, M. Patrikakis, A. Pombo, X. Y. Qin, S. Roy, H. Sato, S. Savvi, A. Saxena, A. Schwegmann, D. Sugiyama, R. Swoboda, H. Tanaka, A. Tomoiu, L. N. Winteringham, E. Wolvetang, C. Yanagi-Mizuochi, M. Yoneda, S. Zabierowski, P. Zhang, I. Abugessaisa, N. Bertin, A. D. Diehl, S. Fukuda, M. Furuno, J. Harshbarger, A. Hasegawa, F. Hori, S. Ishikawa-Kato, Y. Ishizu, M. Itoh, T. Kawashima, M. Kojima, N. Kondo, M. Lizio, T. F. Meehan, C. J. Mungall, M. Murata, H. Nishiyori-Sueki, S. Sahin, S. Nagao-Sato, J. Severin, M. J. de Hoon, J. Kawai, T. Kasukawa, T. Lassmann, H. Suzuki, H. Kawaji, K. M. Summers, C. Wells, F. Consortium, D. A. Hume, A. R. Forrest, A. Sandelin, P. Carninci, Y. Hayashizaki, Transcribed enhancers lead waves of coordinated transcription in transitioning mammalian cells. Science 347, 1010–1014 (2015).

32. B. Langmead, S. L. Salzberg, Fast gapped-read alignment with Bowtie 2. Nat Methods 9, 357–359 (2012).

33. P. Danecek, J. K. Bonfield, J. Liddle, J. Marshall, V. Ohan, M. O. Pollard, A. Whitwham, T. Keane, S. A. McCarthy, R. M. Davies, H. Li, Twelve years of SAMtools and BCFtools. Gigascience 10, (2021).

34. H. M. Amemiya, A. Kundaje, A. P. Boyle, The ENCODE Blacklist: Identification of Problematic Regions of the Genome. Sci Rep 9, 9354 (2019).

35. F. Ramirez, D. P. Ryan, B. Gruning, V. Bhardwaj, F. Kilpert, A. S. Richter, S. Heyne, F. Dundar, T. Manke, deepTools2: a next generation web server for deep-sequencing data analysis. Nucleic Acids Res 44, W160–165 (2016).

36. J. T. Robinson, H. Thorvaldsdottir, W. Winckler, M. Guttman, E. S. Lander, G. Getz, J. P. Mesirov, Integrative genomics viewer. Nat Biotechnol 29, 24–26 (2011).

37. Y. Zhang, T. Liu, C. A. Meyer, J. Eeckhoute, D. S. Johnson, B. E. Bernstein, C. Nusbaum, R. M. Myers, M. Brown, W. Li, X. S. Liu, Model-based analysis of ChIP-Seq (MACS). Genome Biol 9, R137 (2008).

38. A. R. Quinlan, I. M. Hall, BEDTools: a flexible suite of utilities for comparing genomic features. Bioinformatics 26, 841–842 (2010).

39. C. S. Ross-Innes, R. Stark, A. E. Teschendorff, K. A. Holmes, H. R. Ali, M. J. Dunning, G. D. Brown, O. Gojis, I. O. Ellis, A. R. Green, S. Ali, S. F. Chin, C. Palmieri, C. Caldas, J. S. Carroll, Differential oestrogen receptor binding is associated with clinical outcome in breast cancer. Nature 481, 389–393 (2012).

40. G. Yu, L. G. Wang, Q. Y. He, ChIPseeker: an R/Bioconductor package for ChIP peak annotation, comparison and visualization. Bioinformatics 31, 2382–2383 (2015).

41. A. Dobin, C. A. Davis, F. Schlesinger, J. Drenkow, C. Zaleski, S. Jha, P. Batut, M. Chaisson, T. R. Gingeras, STAR: ultrafast universal RNA-seq aligner. Bioinformatics 29, 15–21 (2013).

42. S. Anders, P. T. Pyl, W. Huber, HTSeq--a Python framework to work with high-throughput sequencing data. Bioinformatics 31, 166–169 (2015).

43. M. I. Love, W. Huber, S. Anders, Moderated estimation of fold change and dispersion for RNA-seq data with DESeq2. Genome Biol 15, 550 (2014).

44. A. Subramanian, P. Tamayo, V. K. Mootha, S. Mukherjee, B. L. Ebert, M. A. Gillette, A. Paulovich, S. L. Pomeroy, T. R. Golub, E. S. Lander, J. P. Mesirov, Gene set enrichment analysis: a knowledge-based approach for interpreting genome-wide expression profiles. Proc Natl Acad Sci U S A 102, 15545–15550 (2005).

45. D. W. Barnett, E. K. Garrison, A. R. Quinlan, M. P. Stromberg, G. T. Marth, BamTools: a C++ API and toolkit for analyzing and managing BAM files. Bioinformatics 27, 1691–1692 (2011).

46. N. Servant, N. Varoquaux, B. R. Lajoie, E. Viara, C. J. Chen, J. P. Vert, E. Heard, J. Dekker, E. Barillot, HiC-Pro: an optimized and flexible pipeline for Hi-C data processing. Genome Biol 16, 259 (2015).

47. S. Bhattacharyya, V. Chandra, P. Vijayanand, F. Ay, Identification of significant chromatin contacts from HiChIP data by FitHiChIP. Nat Commun 10, 4221 (2019).

48. T. Chu, Z. Wang, S. P. Chou, C. G. Danko, Discovering Transcriptional Regulatory Elements From Run-On and Sequencing Data Using the Web-Based dREG Gateway. Curr Protoc Bioinformatics 66, e70 (2019).

49. S. Heinz, C. Benner, N. Spann, E. Bertolino, Y. C. Lin, P. Laslo, J. X. Cheng, C. Murre, H. Singh, C. K. Glass, Simple combinations of lineage-determining transcription factors prime cis-regulatory elements required for macrophage and B cell identities. Mol Cell 38, 576–589 (2010).

50. G. X. Zheng, J. M. Terry, P. Belgrader, P. Ryvkin, Z. W. Bent, R. Wilson, S. B. Ziraldo, T. D. Wheeler, G. P. McDermott, J. Zhu, M. T. Gregory, J. Shuga, L. Montesclaros, J. G. Underwood, D. A. Masquelier, S. Y. Nishimura, M. Schnall-Levin, P. W. Wyatt, C. M. Hindson, R. Bharadwaj, A. Wong, K. D. Ness, L. W. Beppu, H. J. Deeg, C. McFarland, K. R. Loeb, W. J. Valente, N. G. Ericson, E. A. Stevens, J. P. Radich, T. S. Mikkelsen, B. J. Hindson, J. H. Bielas, Massively parallel digital transcriptional profiling of single cells. Nat Commun 8, 14049 (2017).

51. M. Chan-Seng-Yue, J. C. Kim, G. W. Wilson, K. Ng, E. F. Figueroa, G. M. O’Kane, A. A. Connor, R. E. Denroche, R. C. Grant, J. McLeod, J. M. Wilson, G. H. Jang, A. Zhang, A. Dodd, S. B. Liang, A. Borgida, D. Chadwick, S. Kalimuthu, I. Lungu, J. M. S. Bartlett, P. M. Krzyzanowski, V. Sandhu, H. Tiriac, F. E. M. Froeling, J. M. Karasinska, J. T. Topham, D. J. Renouf, D. F. Schaeffer, S. J. M. Jones, M. A. Marra, J. Laskin, R. Chetty, L. D. Stein, G. Zogopoulos, B. Haibe-Kains, P. J. Campbell, D. A. Tuveson, J. J. Knox, S. E. Fischer, S. Gallinger, F. Notta, Transcription phenotypes of pancreatic cancer are driven by genomic events during tumor evolution. Nat Genet 52, 231–240 (2020).

