## Supplementary Figure Legends for "KLF5 controls subtype-independent highly interactive enhancers in pancreatic cancer to regulate cell survival"

**fig. S1. Identification of subtype-independent highly interactive enhancers**.

(**A**) Ranked contact counts in AsPC1 (left) and L3.6pl (right) cells from H3K4me3-anchored HiChIP data. Red line depicts the inflection point, approximately 5 for each cell line. (**B**) Heatmap and aggregate plots of chromatin accessibility and H3K27ac in multiple classical and basal-like A cell lines on subtype-independent highly interactive enhancers. Ranked descending on ATAC-seq signal in AsPC1 cells. (**C**) HOMER motif analysis results on the top motifs enriched in subtype-independent highly interactive enhancers. (**D**) Correlation between *KLF5* gene expression (x-axis) and *KLF5* gene effect from CRISPR-dependency data from DepMap.

**fig. S2. KLF5 primarily binds distal regulatory elements.**

(**A**) ChIPseeker analysis showing genome-wide annotation of KLF5 peaks in AsPC1, HPAFII, BxPC3 and L3.6pl cell lines. (**B**) Heatmap of KLF5 binding sites that are unique to each cell line, common (present in all four cell lines) or mixed (present in two or more but not all four cell lines). Ranked descending on KLF5 signal in L3.6pl cells. (**C**) HOMER motif analysis results on the regions described in (B).

**fig. S3. Transcriptomic analysis upon KLF5 depletion.**

(**A**) Representative Western blot analysis of KLF5 after 24 h KLF5 knockdown in AsPC1, HPAFII, BxPC3 and L3.6pl cells. HSC70 was used as a loading control. (**B**) Gene Set Enrichment Analysis (GSEA) after 24 h KLF5 knockdown in AsPC1, HPAFII, BxPC3 and L3.6pl cells. Normalized enrichment score is on x-axis and false discovery rate on y-axis. Common significantly downregulated pathways are displayed on the table. n = 3 biological replicates.

**fig. S4. KLF5 does not alter subtype identity in pancreatic cancer.**

(**A**) GSEA of classical signature (signatures 1 and 6) or basal-like A signature (signature 2) from Chan-Seng-Yue et al. in AsPC1 and HPAFII or BxPC3 and L3.6pl cells, respectively. (**B**) Heatmap of KLF5 at KLF5 unique and KLF5-GATA6 or KLF5-∆Np63 common regions in AsPC1 or L3.6pl cells, respectively.

**fig. S5. KLF5 depletion decreases cell viability.**

(**A**) Representative crystal violet staining after 72 h KLF5 knockdown in AsPC1, HPAFII, BxPC3, L3.6pl and immortalized normal human pancreatic ductal epithelial cell line (HPDEC). (**B**) Quantification of cell confluency of the conditions described in (A). n = 3 biological replicates. (**C**) Representative Western blot analysis of KLF5 after 72 h KLF5 knockdown. HSC70 was used as a loading control.

**fig. S6. Generation of KLF5 degron cell lines.**

(**A**) Schematic depicting genotyping approach for the endogenous knock-in of GFP-FKBP12^F36V^ into the N-terminus of the *KLF5* locus. (**B**) Representative genotyping PCR for N-terminus knockin of *KLF5*. WT band (154 bp) and knockin band (1201 bp) show homozygous knockin in L3.6pl, BxPC3 and AsPC1 cell lines. (**C**) Representative Western blot of KLF5 in clonal knockin cell lines. Observed endogenous KLF5 molecular weight (~55 kDa), EGFP (~26 kDa) and FKBP12^F36V^ (~12 kDa). HSC70 was used as a loading control.

**fig. S7. Effect of KLF5 degradation on H3K4me3 and ATAC levels.**

(**A and B**) Differential binding analysis of genome-wide H3K4me3 ChIP-seq (A) or chromatin accessibility (B) in knockin L3.6pl cells upon 1, 4 and 24 h dTAG treatment. n = 3 biological replicates. (**C**) Overlap of differentially bound H3K27ac and open regions in knockin L3.6pl cells upon 1, 4 and 24 h dTAG treatment.

**fig. S8. Early KLF5-dependent genes connect with multiple KLF5 peaks.**

(**A**) Heatmap shows the raw z-score values generated on vst transformed normalized counts from RNA-seq after 4, 24 and 48 h dTAG treatment in knockin L3.6pl cells. n = 3 biological replicates. (**B**) Heatmap of raw z-score values generated on vst transformed normalized counts from RNA-seq upon 24 h KLF5 knockdown in parental L3.6pl cells ranked on genes in (A). n = 3 biological replicates. (**C**) Gene Set Enrichment Analysis after 4 h dTAG treatment in knockin L3.6pl cells. Normalized enrichment score is on x-axis and false discovery rate on y-axis. Significantly downregulated genes overlap with pathways regulated upon KLF5 knockdown in parental L3.6pl cells. (**D**) Cumulative percentage plots of the number of KLF5 peaks that interact with H3K4me3 marked early (4 h) or late (24 h) KLF5-dependent genes (Log2FC < -0.5, FDR < 0.05). 139 (4 h) and 179 (24 h) genes were included in the analysis. Mann-Whitney test (Wilcoxon rank-sum test), **** p < 0.0001.

**fig. S9. Early KLF5-dependent and highly connected KLF5 regions are transcribed.**

(**A**) Integrated genome viewer tracks of H3K27ac and KLF5 at the *BCL2L1* locus in AsPC1, HPAFII, BxPC3 and L3.6pl cell lines. (**B**) qPCR of *BCL2L1* upon 48 h Zim3-dCas9 simultaneously targeting all three marked enhancers in BxPC3 and HPAFII cells. *ACTB* was used to normalize gene expression. n = 3 biological replicates. (**C to E**) Knockin L3.6pl (C), BxPC3 (D) and AsPC1 (E) cells were plated overnight and treated with 250 nM dTAGV1-TFA and monitored for confluency. n = 6 biological replicates. Quantification of confluency over time.

**fig. S10. KLF5 and Bcl-xL are co-expressed in patients.**

(**A**) Pearson correlation of *KLF5* and *BCL2L1* gene expression in patient single-cell RNA-seq data. (**B**) Representative multiplex immunofluorescence images validating KLF5 and Bcl-xL co-expression in patient samples independent of subtype-identity. Scale bar represents 50 µm.

**fig. S11. KLF5 regulates cell viability partially through apoptosis.**

(**A to D**) BxPC3 and HPAFII cells were treated with KLF5 or non-targeting siRNA mix, incubated for 4 h and media was changed with DMSO or 10 µM Q-VD-OPh with live cell annexin V reagent (1:1,500) and monitored via live cell imaging for proliferation and red calibrated units (RCU). n = 3 biological replicates. (A and C) Quantification of confluency over time. Unpaired Student’s t-test on the area under the curve (AUC), **** p < 0.0001. (B and D) Fold change (siKLF5 vs siNT5) of RCU per % confluency. (**E**) Lentiviral Zim3-dCas9 BxPC3 cells were treated with a pool targeting downstream enhancers of *BCL2L1* or tracrRNA alone, incubated for 4 h and monitored via live cell imaging for proliferation. n = 3 biological replicates. Quantification of confluency over time. Unpaired Student’s t-test on the AUC, **** p < 0.0001. (**F and G**) BxPC3 cells were transfected with either empty vector backbone (pcDNA3.1) or containing Bcl-xL-mCherry for 24 h and subsequently transfected with KLF5 or non-targeting siRNA mix, incubated for 4 h and media was changed with live cell caspase 3/7 reagent (1:1,000) and monitored via live cell imaging for proliferation and GCU and RCU. n = 3 biological replicates. (F) Representative Western blot analysis of KLF5 and Bcl-xL confirming knockdown and overexpression, respectively. HSC70 was used as a loading control. (G) Fraction of cells with only caspase3/7 reagent (GCU) or caspase 3/7 reagent and mCherry (GCU + RCU). (**H and I**) Knockin L3.6pl cells were plated overnight and treated with indicated reagents and monitored for confluency. n = 6 biological replicates. (H) Quantification of confluency over time. (I) Quantification of Bliss synergy score from (H).
