## Supplementary Figures S1-S11 for "KLF5 controls subtype-independent highly interactive enhancers in pancreatic cancer to regulate cell survival"

**Figure S1**

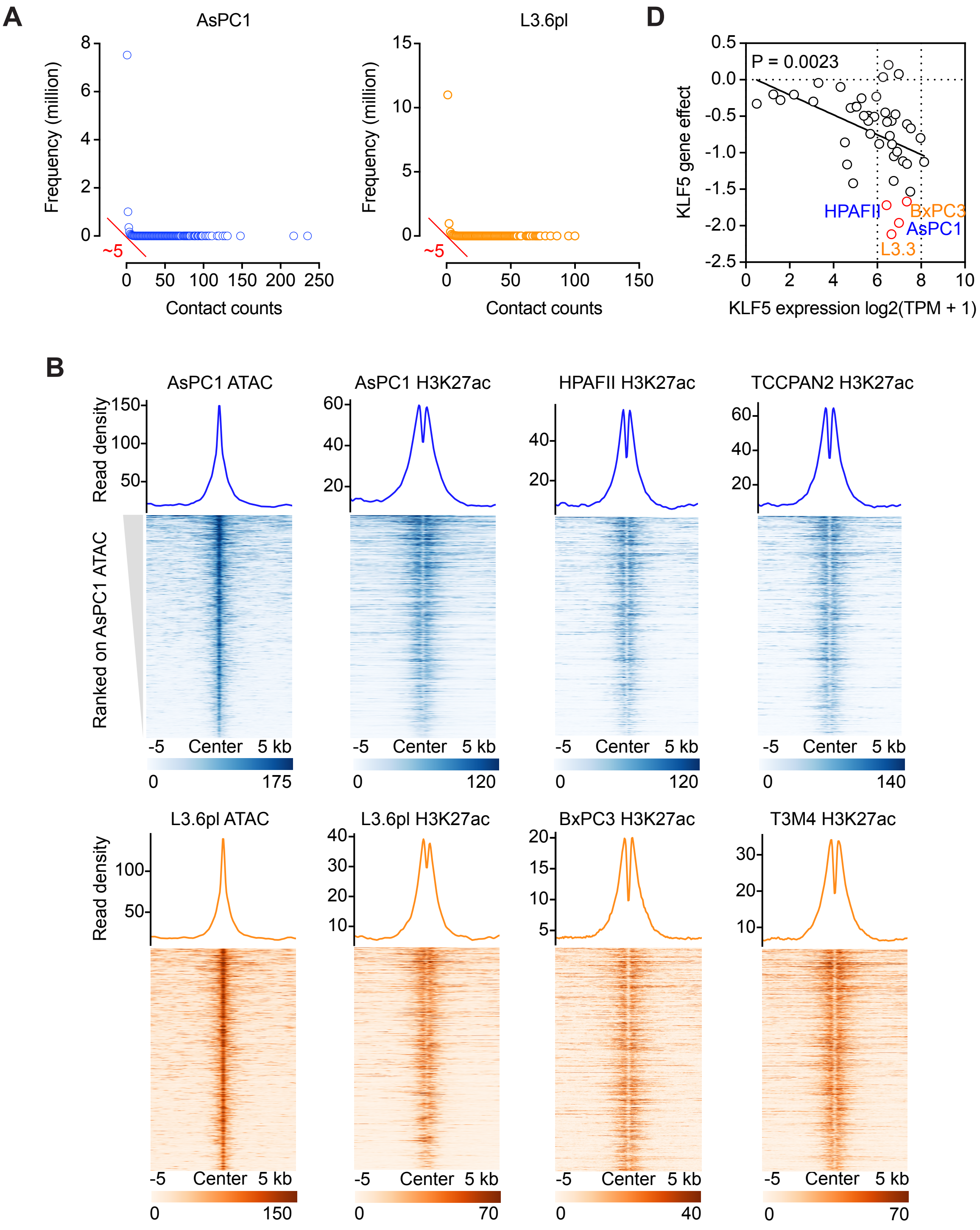


**C**

HOMER motif analysis, P-val ≤ 1e-50 and target sequences with motif ≥ 25%

| Name | Motif | P-val | % of target seq with motif | Name | Motif | P-val | % of target seq with motif |
|---|---|---|---|---|---|---|---|
| FRA1 | 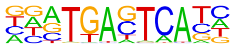 | 1e-330 | 36.18 | BATF | 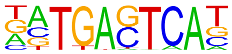 | 1e-308 | 37.50 |
| FOS | 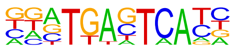 | 1e-324 | 36.79 | JUN-AP1 | 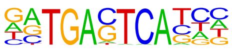 | 1e-293 | 26.03 |
| FRA2 | 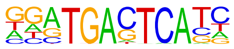 | 1e-320 | 34.12 | AP-1 | 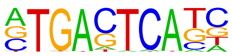 | 1e-288 | 38.47 |
| FOSL2 | 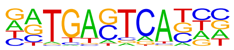 | 1e-318 | 30.23 | ETV1 | 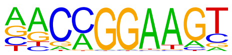 | 1e-52 | 31.45 |
| ATF3 | 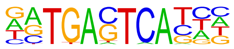 | 1e-315 | 38.23 | KLF5 | 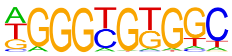 | 1e-50 | 39.31 |
| JUNB | 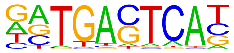 | 1e-312 | 35.50 | | | | |

#### Figure S2

**A**

**B**

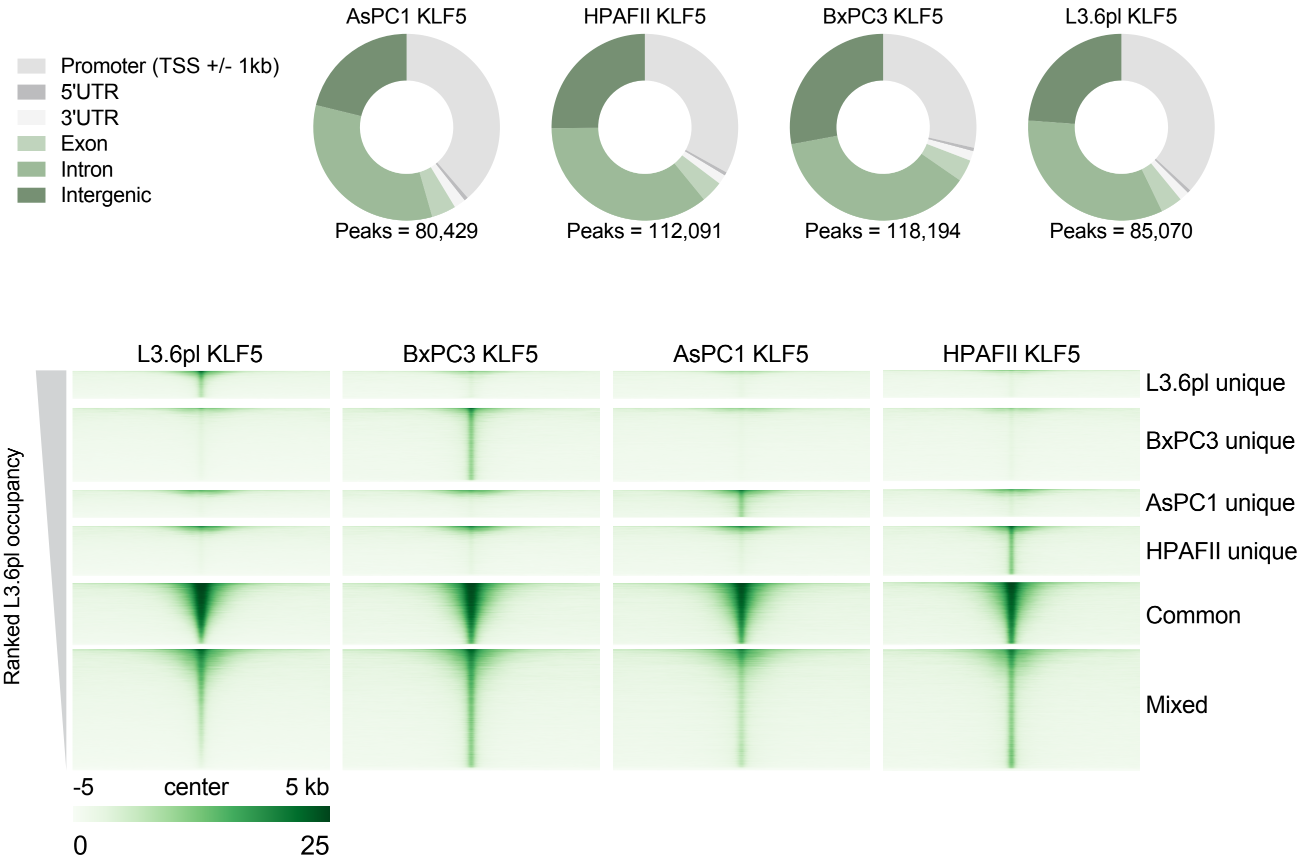


**C**

| | KLF5 *de novo* motif | P-val | Rank | KLF5 known motif | P-val | Rank |
|---|---|---|---|---|---|---|
| AsPC1 unique | KLF5 motif not detected | | | 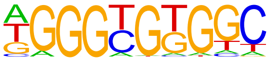 | 1 x -10$^{919}$ | #1 |
| HPAFII unique | 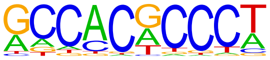 | 1 x -10$^{2255}$ | #1 | | 1 x -10$^{2119}$ | #2 |
| BxPC3 unique | 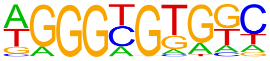 | 1 x -10$^{2775}$ | #1 | | 1 x -10$^{2634}$ | #1 |
| L3.6pl unique | KLF5 motif not detected | | | | 1 x -10$^{433}$ | #3 |
| Common | 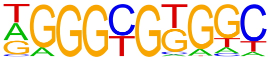 | 1 x -10$^{4067}$ | #1 | | 1 x -10$^{3824}$ | #1 |
| Mixed | 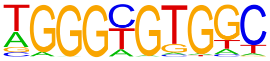 | 1 x -10$^{6320}$ | #1 | | 1 x -10$^{6502}$ | #1 |

#### Figure S3

**A**

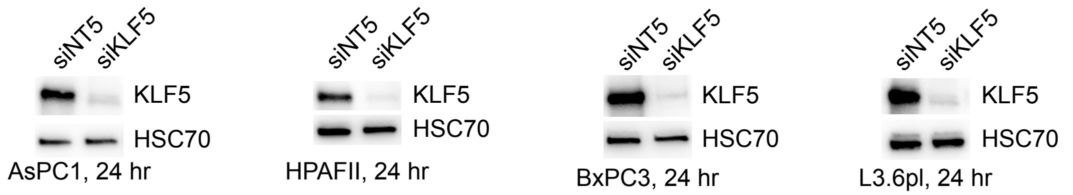


**B**

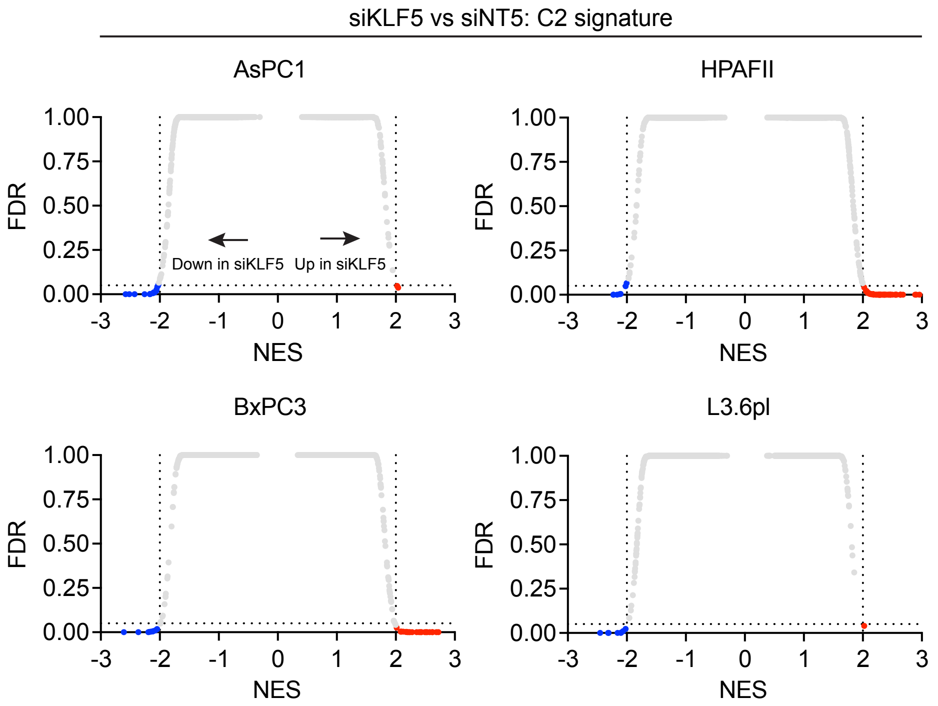


| Common | AsPC1 | HPAFII | BxPC3 | L3.6pl |
|---|---|---|---|---|
| Anderson cholangiocarcinoma class 2 | -2.57 | -2.57 | -2.00 | -2.45 |
| Reactome keratinization | -2.11 | -2.07 | -2.09 | -2.15 |
| Reactome of the cornified envelope | -2.16 | -2.15 | -2.22 | -2.09 |
| Cromer metastasis down | -2.51 | -2.02 | -2.36 | -2.02 |

**C**

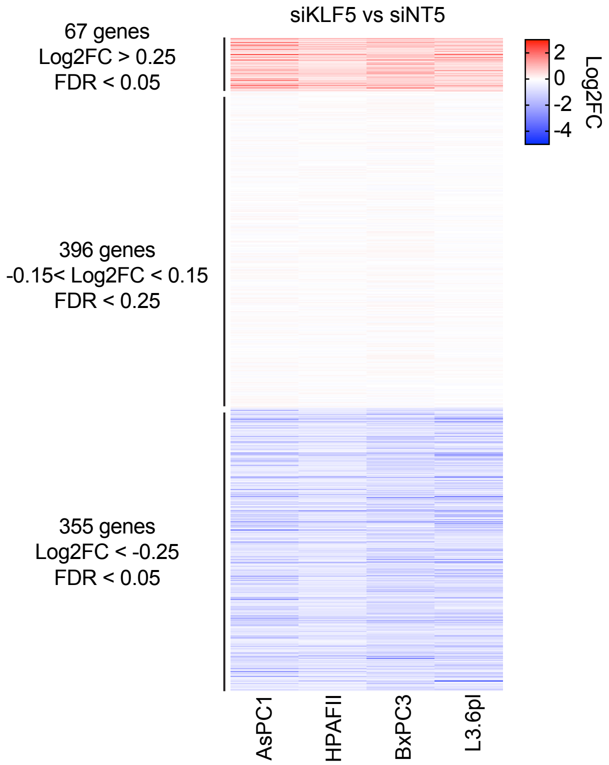

#### Figure S4

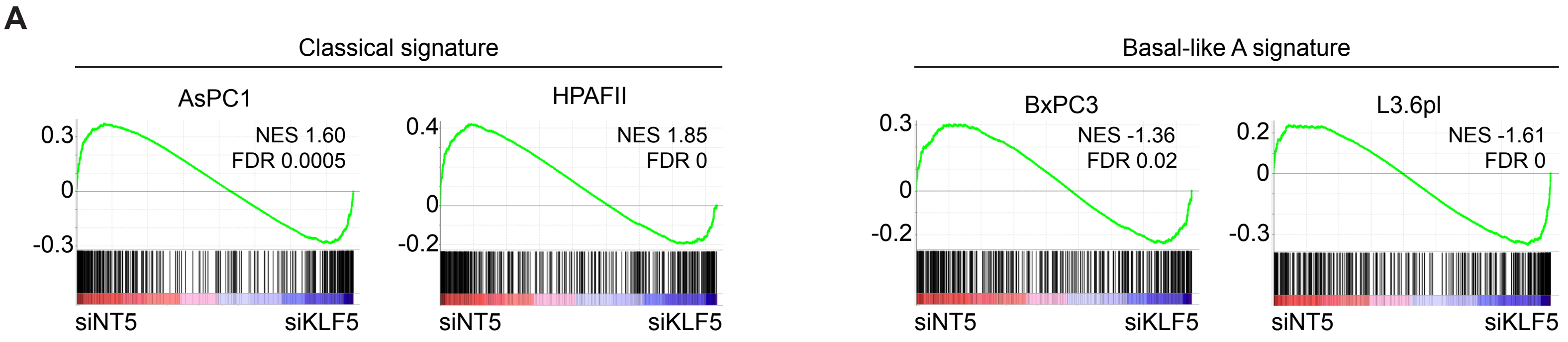
A
Classical signature
AsPC1
0.3
0
-0.3
NES 1.60
FDR 0.0005
siNT5
siKLF5
HPAFII
0.4
0
-0.2
NES 1.85
FDR 0
siNT5
siKLF5
Basal-like A signature
BxPC3
0.3
0
-0.2
NES -1.36
FDR 0.02
siNT5
siKLF5
L3.6pl
0.2
0
-0.3
NES -1.61
FDR 0
siNT5
siKLF5

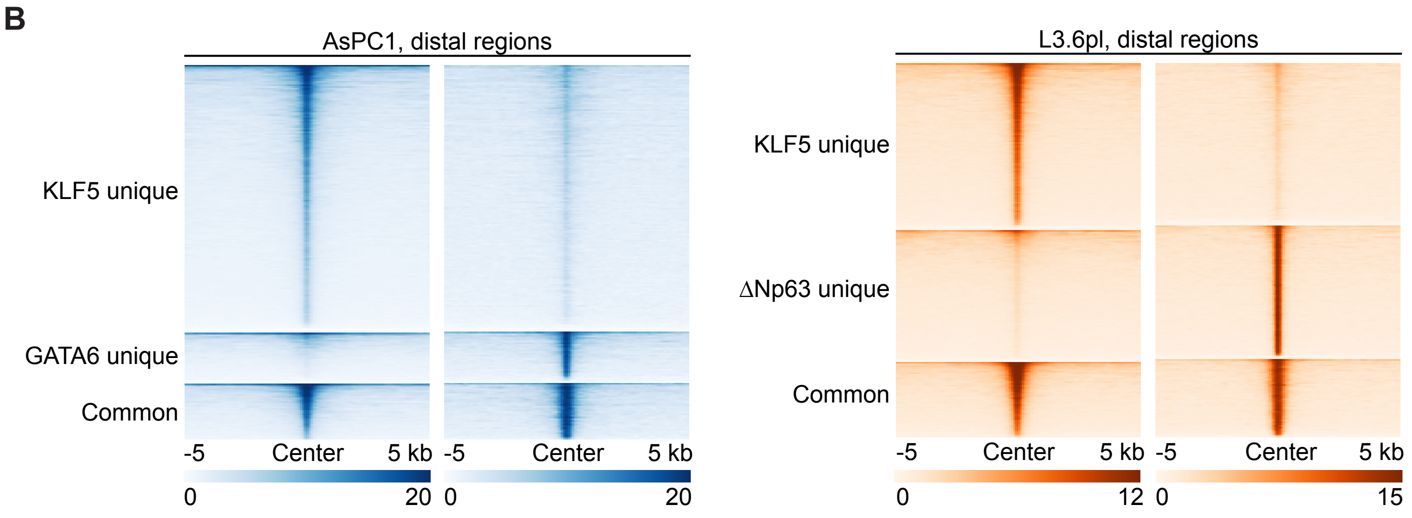
B
AsPC1, distal regions
KLF5 unique
GATA6 unique
Common
-5
Center
5 kb
-5
Center
5 kb
0
20
0
20
L3.6pl, distal regions
KLF5 unique
ΔNp63 unique
Common
-5
Center
5 kb
-5
Center
5 kb
0
12
0
15

**Figure S5**

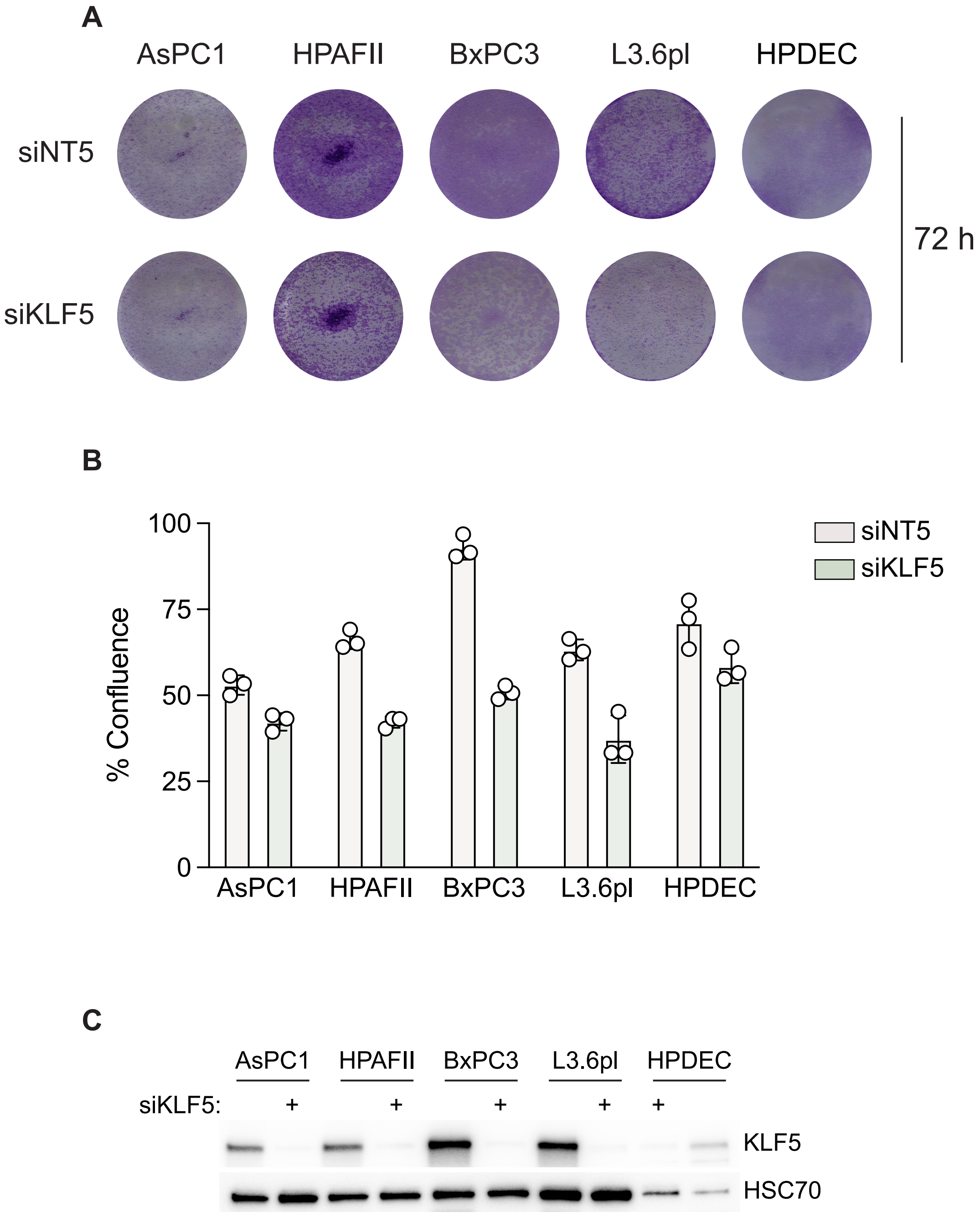
A
AsPC1
HPAFII
BxPC3
L3.6pl
HPDEC
siNT5
siKLF5
72 h
B
100
75
50
25
0
% Confluence
siNT5
siKLF5
AsPC1
HPAFII
BxPC3
L3.6pl
HPDEC
C
AsPC1
HPAFII
BxPC3
L3.6pl
HPDEC
siKLF5:
+
+
+
+
+
KLF5
HSC70

#### Figure S6

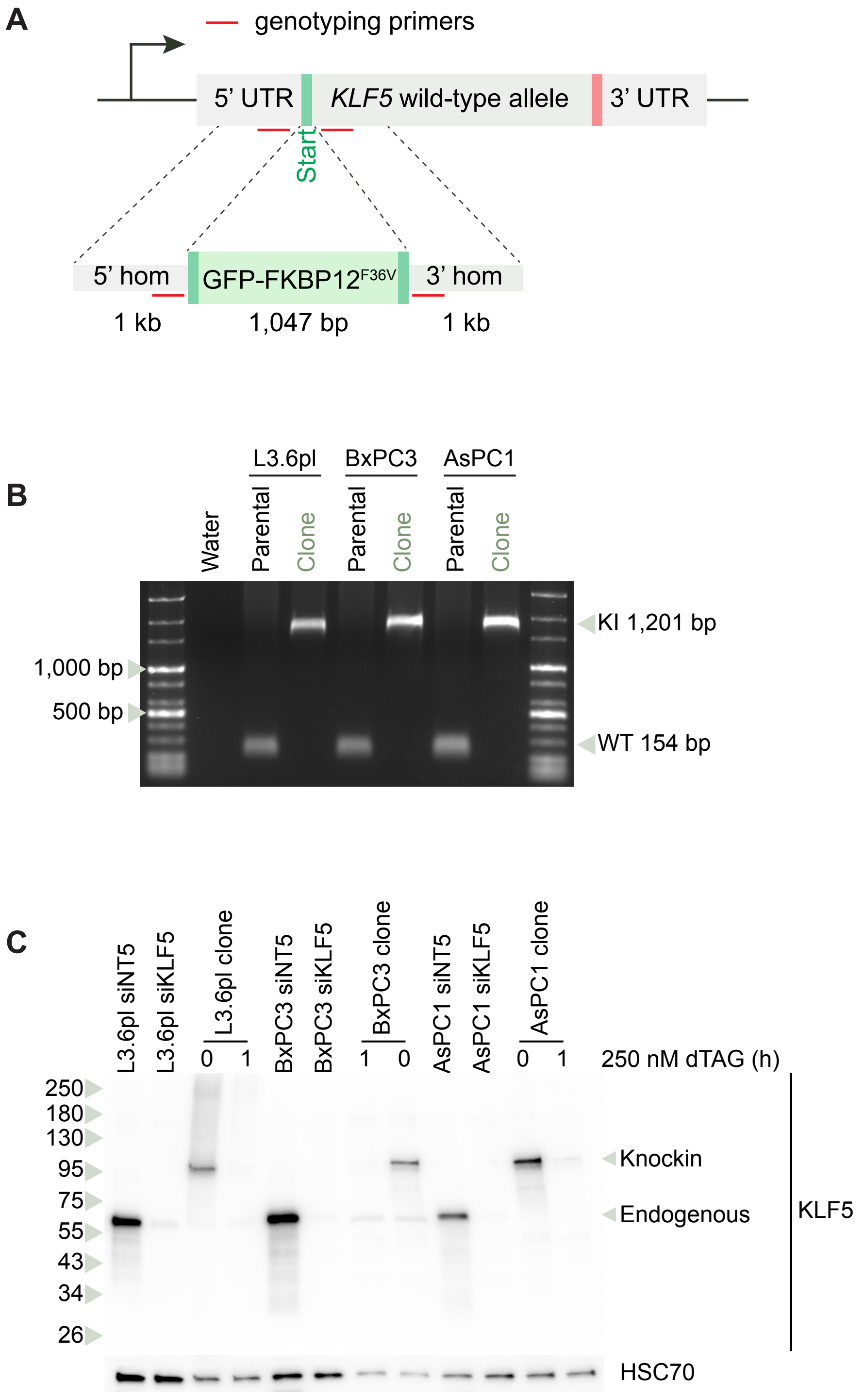
A
genotyping primers
5' UTR
KLF5 wild-type allele
3' UTR
Start
5' hom
GFP-FKBP12F36V
3' hom
1 kb
1,047 bp
1 kb
B
L3.6pl
BxPC3
AsPC1
Water
Parental
Clone
Parental
Clone
Parental
Clone
1,000 bp
500 bp
KI 1,201 bp
WT 154 bp
C
L3.6pl siNT5
L3.6pl siKLF5
L3.6pl clone
BxPC3 siNT5
BxPC3 siKLF5
BxPC3 clone
AsPC1 siNT5
AsPC1 siKLF5
AsPC1 clone
0
1
1
0
0
1
250 nM dTAG (h)
250
180
130
95
75
55
43
34
26
Knockin
Endogenous
KLF5
HSC70

#### Figure S7

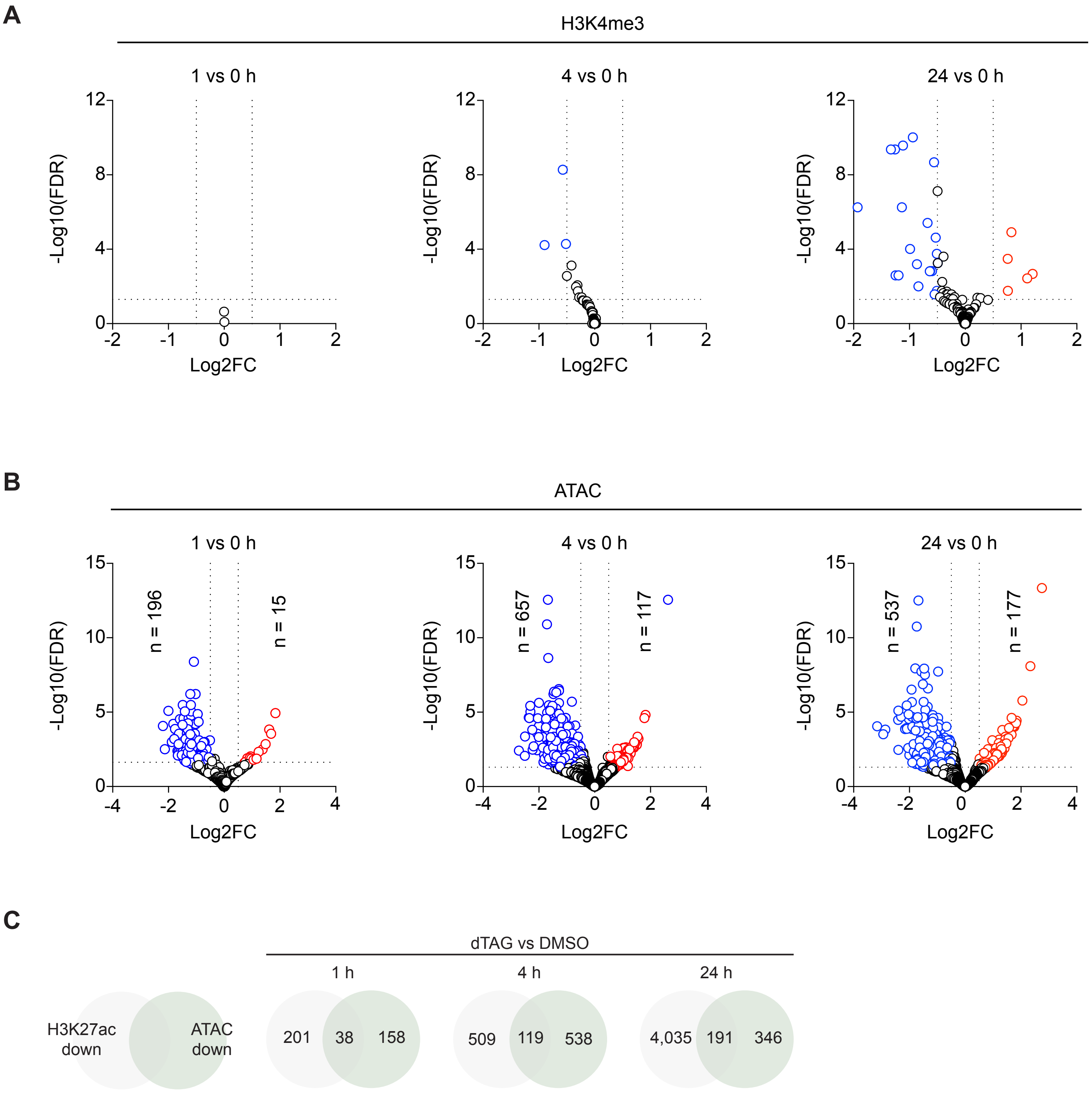
A
H3K4me3
1 vs 0 h
4 vs 0 h
24 vs 0 h
-Log10(FDR)
Log2FC
B
ATAC
1 vs 0 h
4 vs 0 h
24 vs 0 h
n = 196
n = 15
n = 657
n = 117
n = 537
n = 177
-Log10(FDR)
Log2FC
C
dTAG vs DMSO
1 h
4 h
24 h
H3K27ac down
ATAC down
201
38
158
509
119
538
4,035
191
346

#### Figure S8

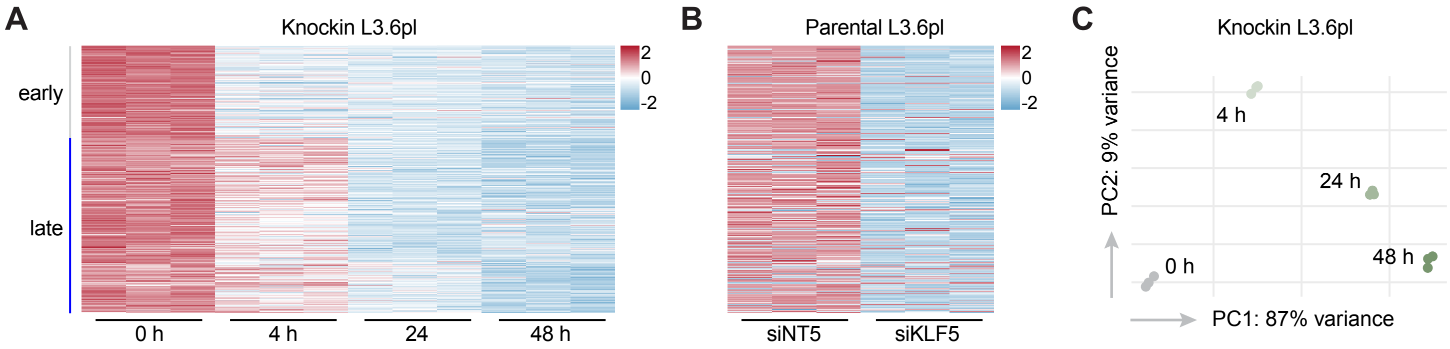


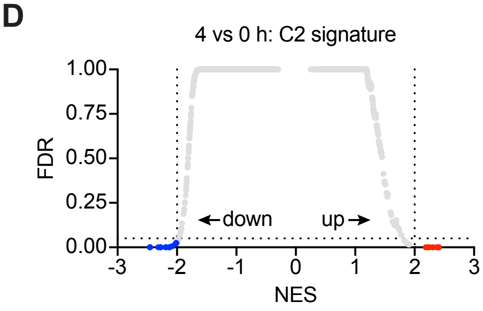


| Common regulated gene sets | | | |
|---|---|---|---|
| Anderson cholangiocarcinoma class 2 | -2.45 | Reactome of the cornified envelope | -2.12 |
| Reactome keratinization | -2.27 | Cromer metastasis down | -2.27 |

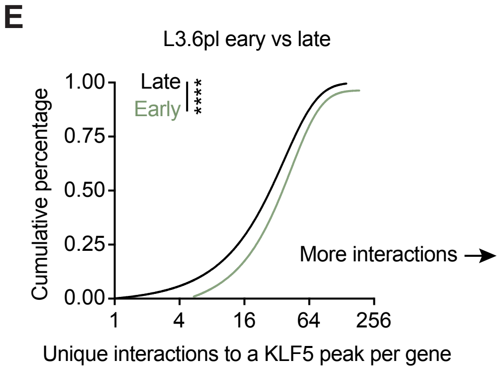

#### Figure S9

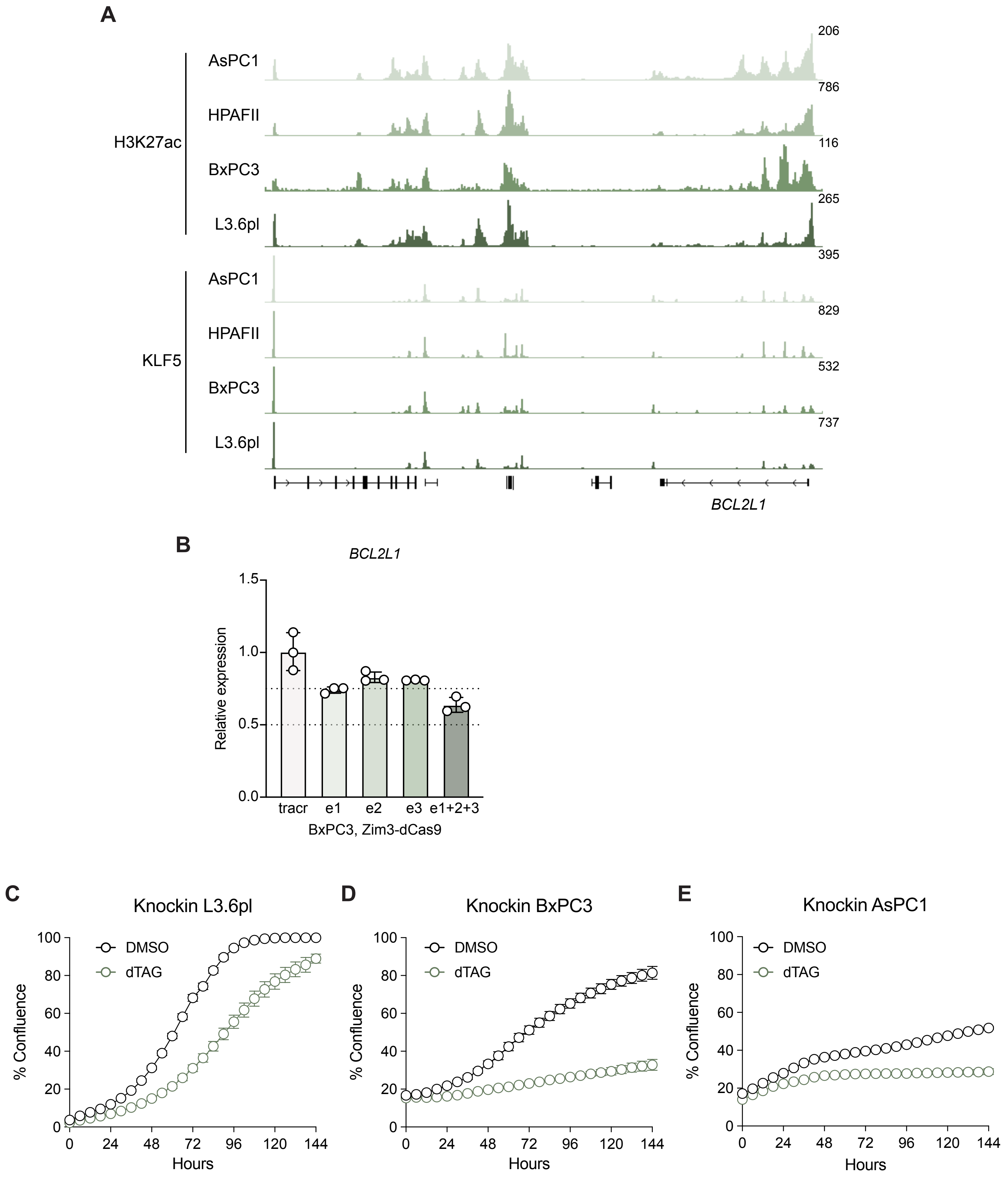
A
H3K27ac
AsPC1
HPAFII
BxPC3
L3.6pl
KLF5
AsPC1
HPAFII
BxPC3
L3.6pl
206
786
116
265
395
829
532
737
BCL2L1
B
BCL2L1
Relative expression
1.5
1.0
0.5
0.0
tracr
e1
e2
e3
e1+2+3
BxPC3, Zim3-dCas9
C
Knockin L3.6pl
D
Knockin BxPC3
E
Knockin AsPC1
DMSO
dTAG
% Confluence
100
80
60
40
20
0
0
24
48
72
96
120
144
Hours

#### Figure S10

A

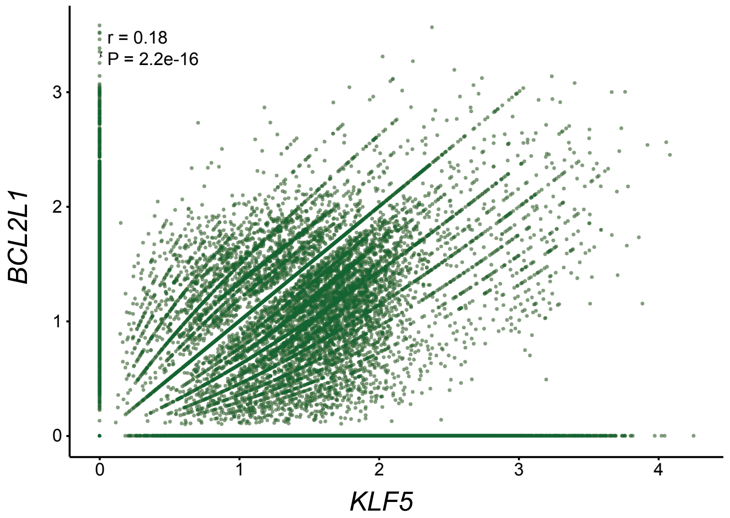


B

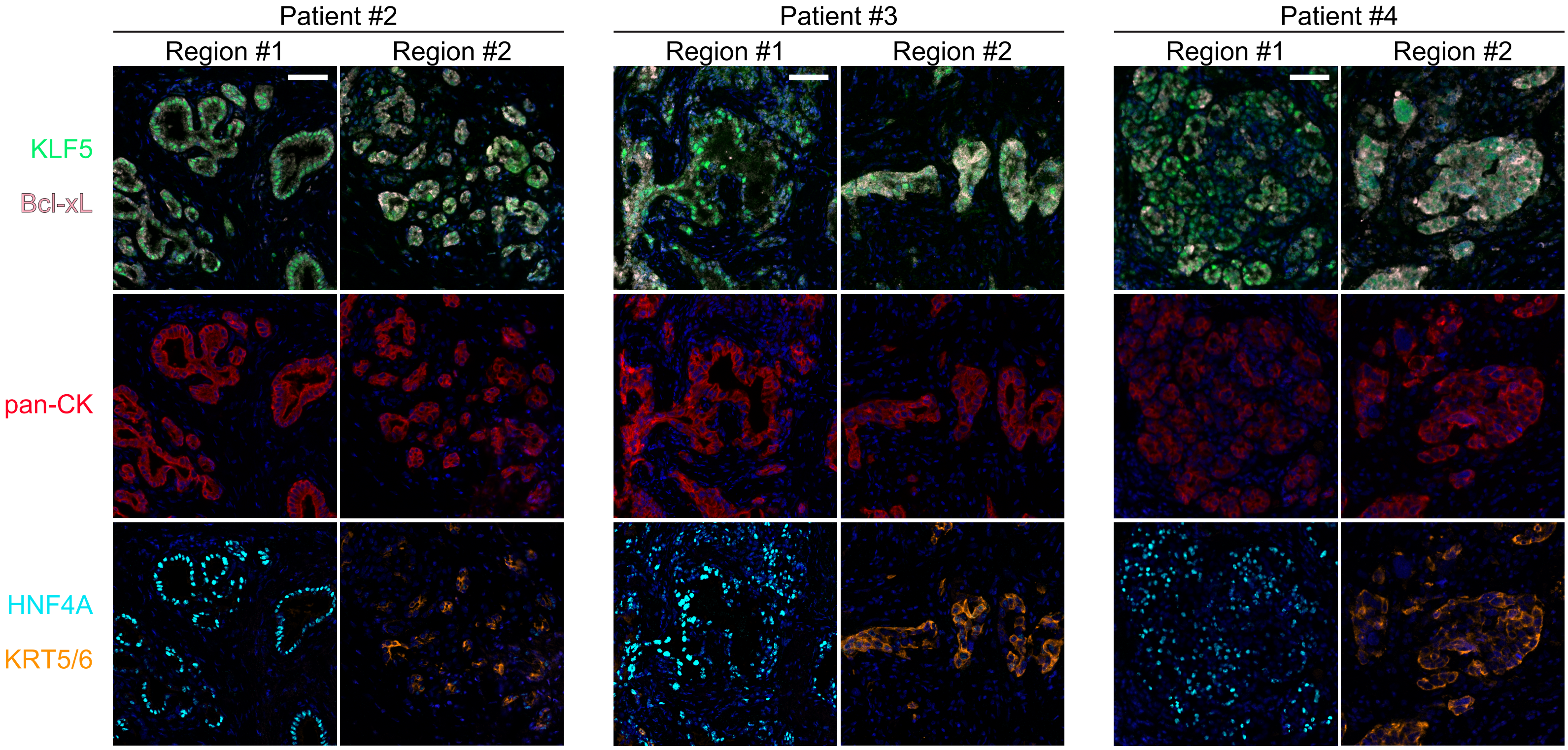

### Figure S11

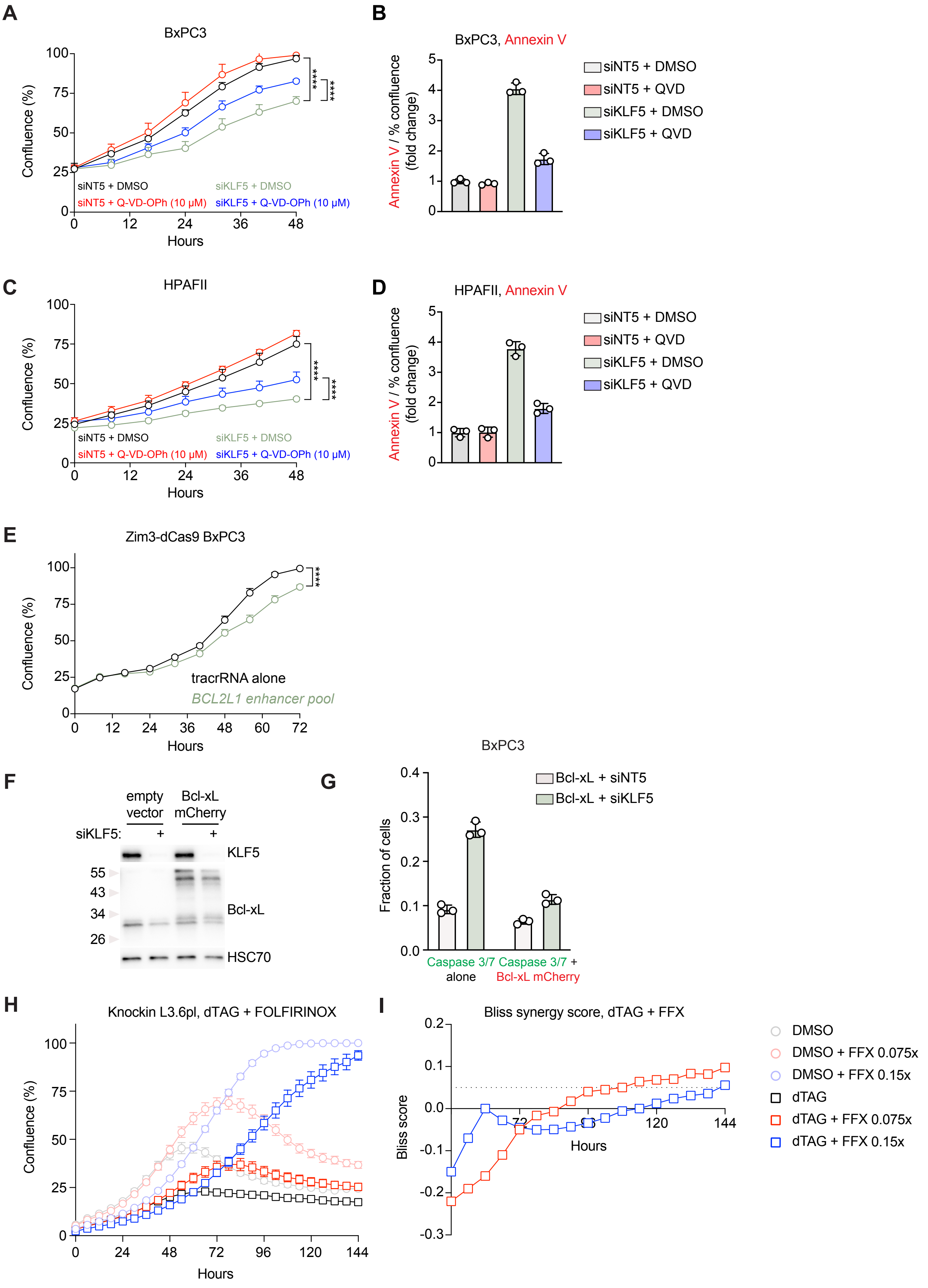

A
BxPC3
Confluence (%)
Hours
siNT5 + DMSO
siKLF5 + DMSO
siNT5 + Q-VD-OPh (10 μM)
siKLF5 + Q-VD-OPh (10 μM)
****
****
B
BxPC3, Annexin V
Annexin V / % confluence (fold change)
siNT5 + DMSO
siNT5 + QVD
siKLF5 + DMSO
siKLF5 + QVD
C
HPAFII
Confluence (%)
Hours
siNT5 + DMSO
siKLF5 + DMSO
siNT5 + Q-VD-OPh (10 μM)
siKLF5 + Q-VD-OPh (10 μM)
D
HPAFII, Annexin V
Annexin V / % confluence (fold change)
siNT5 + DMSO
siNT5 + QVD
siKLF5 + DMSO
siKLF5 + QVD
E
Zim3-dCas9 BxPC3
Confluence (%)
Hours
tracrRNA alone
BCL2L1 enhancer pool
F
empty vector
Bcl-xL mCherry
siKLF5:
KLF5
Bcl-xL
HSC70
55
43
34
26
G
BxPC3
Fraction of cells
Bcl-xL + siNT5
Bcl-xL + siKLF5
Caspase 3/7 alone
Caspase 3/7 + Bcl-xL mCherry
H
Knockin L3.6pl, dTAG + FOLFIRINOX
Confluence (%)
Hours
I
Bliss synergy score, dTAG + FFX
Bliss score
Hours
DMSO
DMSO + FFX 0.075x
DMSO + FFX 0.15x
dTAG
dTAG + FFX 0.075x
dTAG + FFX 0.15x
